# Dual-Action Kinase Inhibitors Influence p38α MAP Kinase Dephosphorylation

**DOI:** 10.1101/2024.05.15.594272

**Authors:** Emily J Stadnicki, Hannes Ludewig, Ramasamy P Kumar, Xicong Wang, Youwei Qiao, Dorothee Kern, Niels Bradshaw

## Abstract

Reversible protein phosphorylation directs essential cellular processes including cell division, cell growth, cell death, inflammation, and differentiation. Because protein phosphorylation drives diverse diseases, kinases and phosphatases have been targets for drug discovery, with some achieving remarkable clinical success. Most protein kinases are activated by phosphorylation of their activation loops, which shifts the conformational equilibrium of the kinase towards the active state. To turn off the kinase, protein phosphatases dephosphorylate these sites, but how the conformation of the dynamic activation loop contributes to dephosphorylation was not known. To answer this, we modulated the activation loop conformational equilibrium of human p38α MAP kinase with existing kinase inhibitors that bind and stabilize specific inactive activation loop conformations. From this, we discovered three inhibitors that increase the rate of dephosphorylation of the activation loop phospho-threonine by the PPM serine/threonine phosphatase WIP1. Hence, these compounds are “dual-action” inhibitors that simultaneously block the active site and stimulate p38α dephosphorylation. Our X-ray crystal structures of phosphorylated p38α bound to the dual-action inhibitors reveal a shared flipped conformation of the activation loop with a fully accessible phospho-threonine. In contrast, our X-ray crystal structure of phosphorylated apo human p38α reveals a different activation loop conformation with an inaccessible phospho-threonine, thereby explaining the increased rate of dephosphorylation upon inhibitor binding. These findings reveal a conformational preference of phosphatases for their targets and suggest a new approach to achieving improved potency and specificity for therapeutic kinase inhibitors.

## Introduction

Reversible protein phosphorylation is widely deployed to control essential cellular physiology including cell division, cell growth, cell death, stress response, inflammation, and differentiation^1,2^. For these purposes, kinases and phosphatases are subject to exquisite regulation, including phosphorylation and dephosphorylation by other kinases and phosphatases, respectively, thus providing cells with an adaptable and highly-interconnected signaling network^3–5^. Because misregulation of these cellular pathways cause diverse diseases, kinases and phosphatases have been targets involved in the most intense efforts for drug-development over the past 25 years^6–9^. Despite the remarkable clinical success of some kinase inhibitors, broader application has been hampered by difficulty achieving specificity due to the highly conserved kinase active site^7^. Targeting phosphatases has been even more challenging due to their lack of a traditionally druggable pocket and the fact that phosphatase activation, rather than inhibition, is advantageous for many therapeutic applications^9^. An attractive but elusive class of compounds would selectively direct phosphatase activity to a particular target protein or phosphorylation site, such as a kinase activation loop. Recently some success has been achieved towards this goal by localizing a phosphatase to its target using heterobifunctional compounds^10,11^, compounds that stabilize phosphatase/adapter complexes^12^, or a transgenic affinity-tagged phosphatase^13^. These proof-of-concept studies have suggested prospective benefits for specificity, potency, and kinetics from phosphatase-driven inhibition. However, the investigated molecules lack drug-like properties and/or relied on transgenic phosphatases for molecular targeting. One example identified a “phosTAC” that recruits the phosphatase to its kinase substrate using an active site competitive EGFR inhibitor tethered to a ligand that recruits a PTPN2-FKBP fusion protein^10^. Similarly, Akt kinase was targeted with heterobifunctional molecules combining active site competitive Akt inhibitors with a ligand to recruit a HALO-PP1 fusion protein or a PP1-activating peptide^14^. A complementary strategy, which we describe here, is to directly target the conformational state of the kinase to increase the rate of dephosphorylation (Fig 1A).

**Figure 1:**
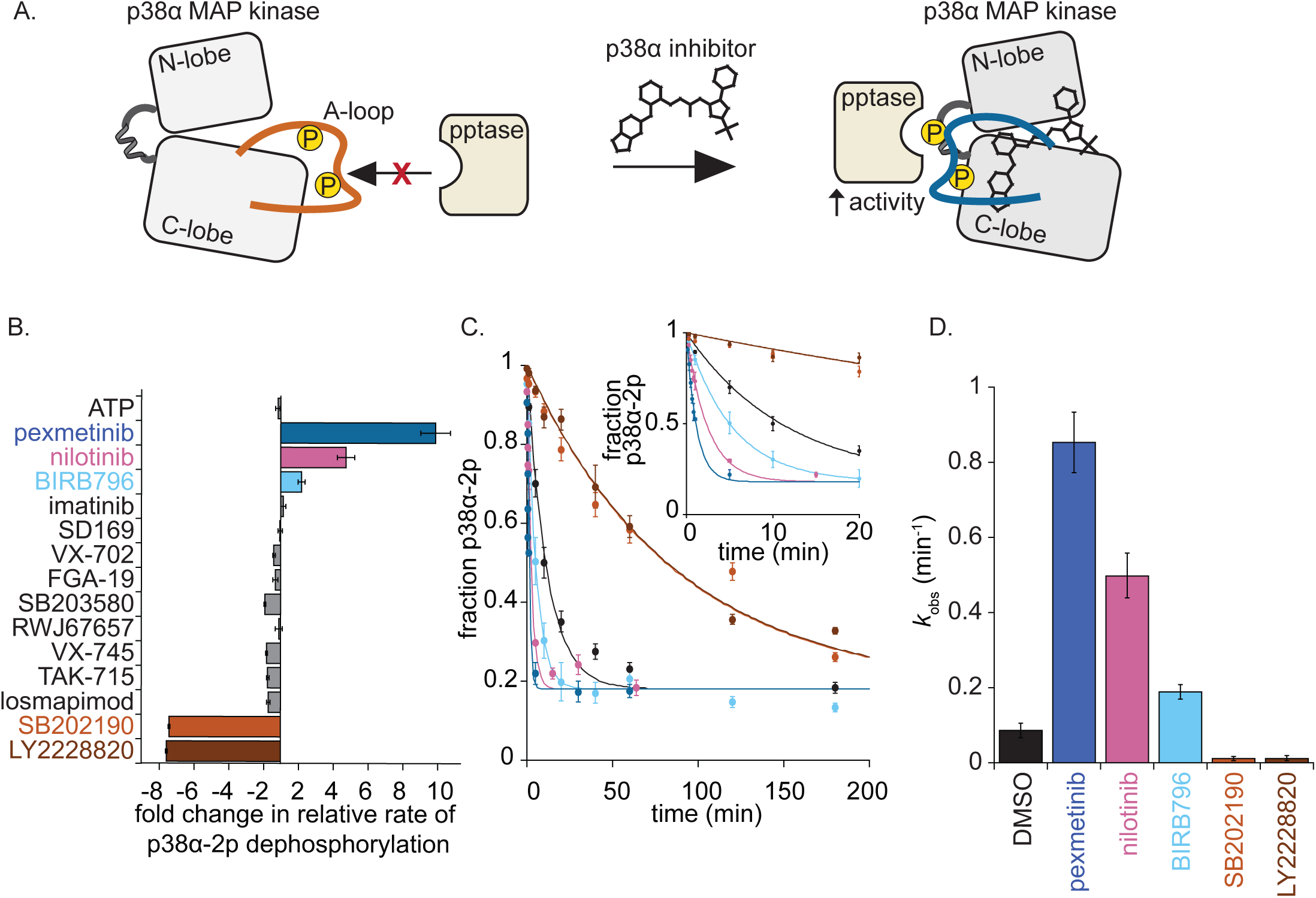
p38α inhibitors modulate dephosphorylation of p38α by WIP1 phosphatase. (A) Schematic depicting p38α inhibitors changing activation loop (A-loop) conformation (apo orange and blue drug bound), leading to a change in phosphatase activity on phosphorylated p38α MAP Kinase. Phosphorylation sites are depicted as yellow spheres on activation loop cartoon. (B) Fold change in single turnover WIP1 phosphatase activity (*k*_obs_ (min^-1^)) in the presence of inhibitors relative to a DMSO treated control (error is the propagated error of the fit). Data for the compounds that caused the largest effects are shown in (C). Observed rates were obtained from single turnover reactions (*k*_obs_ (min^-1^)) of p38α (0.25 µM) dephosphorylation at pT180 by WIP1 (2.5 µM) in the absence or presence of excess p38α inhibitors (1.25 µM). Data are fit to an exponential decay (see methods section) (error on datapoints are ± 1 SD from an n=3). Inset depicts the first 20 minutes of the reaction. (D) *k*_obs_ from panel C shown as bar graphs (*k*_obs_ were averaged from an n=3 ± 1 SD).

The dynamic activation loops of kinases often contain phosphorylation sites that control kinase activity by shifting the conformational ensemble towards states that organize the catalytic center and promote substrate binding. In contrast, kinase inhibitors frequently project into a variable allosteric pocket near the active site, thereby displacing the activation loop and selecting inactive conformations of the kinase^6,8,15^. We postulated that the conformational states adopted by inhibitor bound kinases would impact the rate of dephosphorylation by protein phosphatases, providing an experimental approach to identify kinase conformations that are favorable for dephosphorylation. If this is the case, it suggests a simple mechanism that leverages phosphatases for “dual-action” inhibition, in which the inhibitor simultaneously blocks the active site and directs inactivation of the kinase by dephosphorylation of the activation loop.

We chose the MAP kinase p38α, which is a critical regulatory node for DNA damage response and inflammatory pathways^16–19^, to test whether conformationally selective kinase inhibitors modulate activation loop dephosphorylation. p38α is activated by dual-phosphorylation of its activation loop (p38α-2p) on threonine (pT180) and tyrosine (pY182) residues. T180 phosphorylation stimulates kinase activity more than 1,000-fold and is sufficient for kinase activity in cells, whereas Y182 phosphorylation primarily controls autophosphorylation and contributes two- to ten-fold towards activity^20,21^. In the cell, p38α phosphorylation is controlled by upstream kinases, autophosphorylation, and a suite of protein phosphatases^18^, including serine/threonine phosphatases WIP1^22–24^, PPM1A^25^, and PP2A^26–29^ as well as tyrosine phosphatases (PTPs)^30^, and dual-specificity phosphatases (DUSPs)^22^. Because p38α activation controls cell-death pathways that drive cancer progression and inflammatory responses that cause diseases including myocardial ischemia and neurodegeneration, diverse p38α-specific ATP-competitive inhibitors have been identified and studied in the clinic^31,32^. Many of these inhibitors achieve specificity by extending beyond the ATP binding pocket, which would displace the activation loop from the canonical active conformation, making them ideal candidates to test whether activation loop conformation modulates phosphatase activity.

We identified three conformation-selective p38α inhibitors that stimulate threonine dephosphorylation of the p38α activation loop by WIP1. Our X-ray crystal structures reveal that these compounds favor an activation loop flipped conformation of p38α that presents pT180 for dephosphorylation by WIP1. This demonstrates that the conformation of the activation loop is a critical determinant of the dephosphorylation rate, provides a simple mechanism to promote dephosphorylation of a particularly important regulatory site by a specific phosphatase(s), and provides a roadmap for the development of dual-action kinase inhibitors.

## Results

### Conformation-selective p38α MAP kinase inhibitors modulate threonine dephosphorylation by WIP1

To test the hypothesis that the conformation of the activation loop determines the rate of p38α dephosphorylation, we controlled the activation loop conformational state by binding dual-phosphorylated p38α (p38α-2p) to ATP and a panel of inhibitors (Fig. 1B-D, S1, S2, & S3). We then measured dephosphorylation of the activation loop threonine (pT180) by WIP1, a serine/threonine phosphatase that natively targets pT180 of p38α and has been implicated in oncogenesis^23^. From an initial panel of thirteen inhibitors, we found two related compounds, pexmetinib and BIRB796, that increased the rate of pT180 dephosphorylation 10-fold and 2-fold, respectively. Additionally, we identified two compounds, SB202190 and LY2228820, that each decreased the rate of dephosphorylation 7-fold (Fig. 1B-D, S1A, S2 & S3). To confirm that the compounds act through binding to p38α-2p, we measured WIP1 hydrolysis of the generic substrate fluorescein-diphosphate in the presence of the inhibitors (Fig. S1B) and observed no change in hydrolysis rate, indicating that they do not directly control WIP1 activity. Therefore, we conclude that conformation-selective p38α inhibitors can modulate the rate of WIP1 catalyzed p38α dephosphorylation. Furthermore, we identify two compounds, pexmetinib and BIRB796, that directly inhibit p38α kinase activity and promote p38α inactivation through pT180 dephosphorylation, leading us to classify them as dual-action inhibitors.

### Pexmetinib and BIRB796 present phospho-threonine for dephosphorylation

To determine how WIP1 dephosphorylation of p38α-2p is stimulated by dual-action inhibitors, we solved X-ray crystal structures of human p38α-2p in unliganded- and pexmetinib-bound states (Fig. 2; PDB: 9CJ2 & 9CJ3, respectively & Table S1). In the unliganded form, the two chains of our p38α-2p structure showed slight differences in the closure of the N-lobe and the ordering of the P-loop compared to a previous structure of murine p38α-2p^33^ (PDB: 3PY3). In all three cases, the activation loop is oriented to the right of the active site. While the coordination of the phosphates differs slightly, both phospho-sites are inaccessible to WIP1, which catalyzes dephosphorylation via planar nucleophilic attack of the phosphate by a metal-activated water^34–37^ (Fig. 2A & S4).

**Figure 2:**
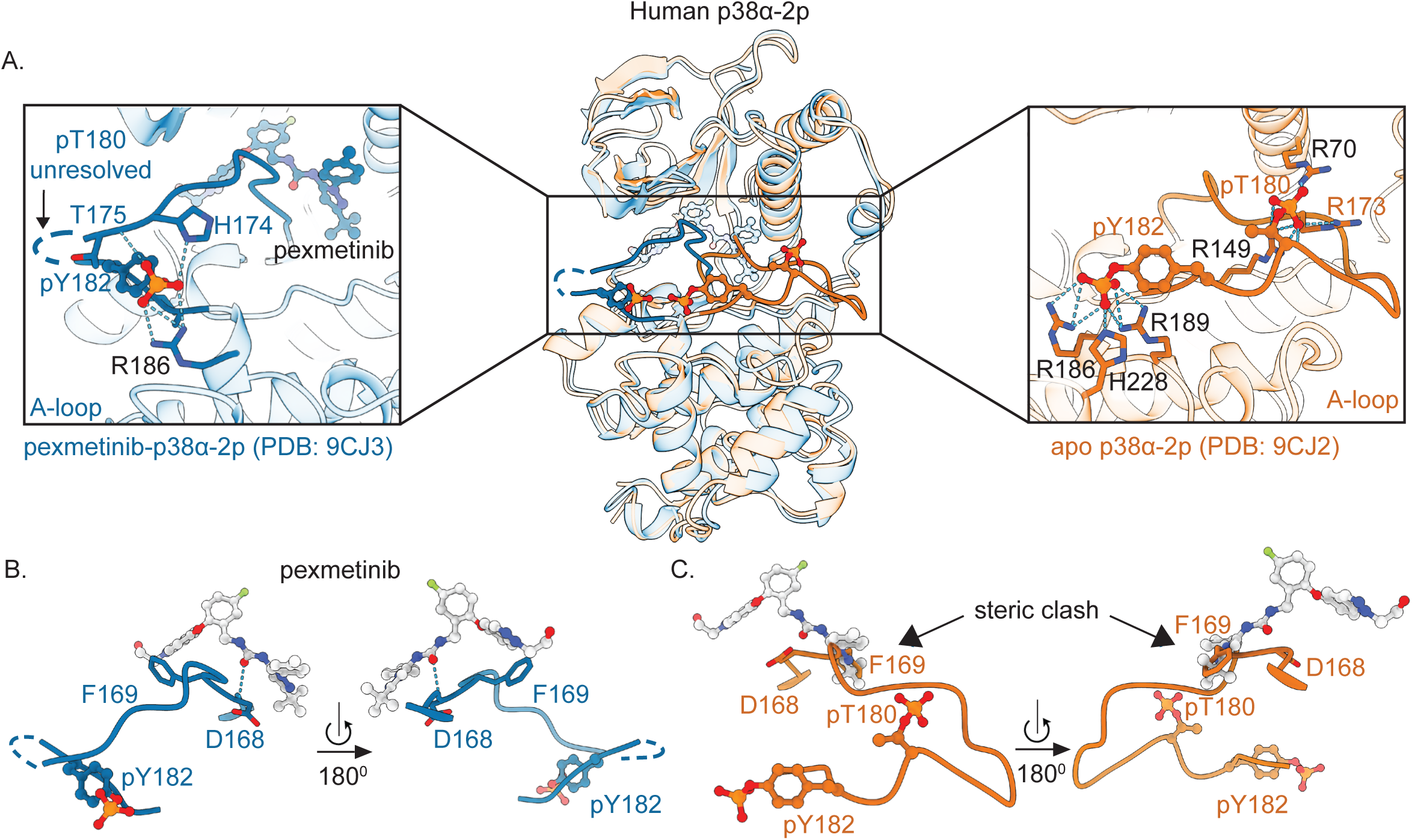
Dual-action p38α inhibitors induce an activation loop conformation that exposes pT180 for dephosphorylation. (A) Overlay of X-ray crystal structure of human apo p38α-2p (orange; PDB: 9CJ2) and pexmetinib bound human p38α-2p (dark blue; PDB: 9CJ3) emphasizing activation loop (A-loop, solid colors) and phosphorylation site rearrangements (shown as sticks). Left zoom shows pexmetinib mediated A-loop conformation and coordination of pY182. Note that pT180 is unresolved, likely due to flexibility. Right zoom shows apo p38α-2p A-loop and coordination of pT180 and pY182. Light blue dotted lines indicate hydrogen bonds to the phosphate group in both zooms. (B) Activation loop of p38α-2p bound pexmetinib (dark blue). Light blue dotted lines indicate hydrogen bonding of pexmetinib to D168 (DFG motif). (C) Activation loop of p38α-2p (orange) overlayed with pexmetinib from the bound structure to emphasize the clash of pexmetinib with F169. In all structures, oxygen, nitrogen, phosphorous and fluorine atoms are colored red, blue, orange and light green, respectively. Crystal contacts are illustrated in Fig. S13.

In our pexmetinib-bound structure of p38α-2p, the activation loop is flipped to the opposite site (Fig. 2A-B). Importantly, this exposes pT180 to solvent, which would render it accessible for a phosphatase to attack. In fact, pT180 and the four preceding amino acids are not resolved in the electron density map, suggestive of flexibility in this region (Fig. 2A-B & S5 A & D). The flipped conformation of the pexmetinib-p38α-2p activation loop is stabilized from the N-terminal end by interactions between the drug and the conserved DFG motif that serves as a hinge for the activation loop (Fig. 2B & C). The carbonyl oxygen of pexmetinib coordinates this interaction through hydrogen bonding with the backbone nitrogen of D168 while the indazole moiety of the drug makes hydrophobic interactions with F169 (Fig. 2A-B). Additionally anchoring the flipped activation loop, the phosphate of pY182 makes hydrogen bonds to the sidechains of R186 and H174, and the backbone of T175 (Fig. 2A). The side chain of R186, which coordinates pY182 in our apo structures, is flipped in murine apo p38α-2p to the position we detected in our human pexmetinib-bound p38α-2p (Fig. 2A), suggesting that coordination of the phosphate is flexible. Our results suggest that pexmetinib promotes p38α dephosphorylation by flipping the activation loop to present pT180 for dephosphorylation by WIP1.

Bolstering our conclusion that activation loop flipping drives dephosphorylation, a similar flip occurs in our structure of p38α-2p bound to the second dual-action inhibitor BIRB796, which stimulated WIP1 dephosphorylation to a lesser extent (Fig. 1B-D, 3A-B, S1A, S5B, & Table S1; PDB: 9CJ4). Like pexmetinib, BIRB796 anchors the activation loop in a flipped conformation through interaction with D168 and F169 from the N-terminal side (Fig. 3A-D, S2, & S5B & E), and pY182 docks in a similar position as in the pexmetinib-bound structure (Fig. 3 & S5D-E). The activation loop is displaced from the kinase-active position by a moiety of BIRB796 that is shared with pexmetinib (Fig. 3B & S2) and clashes with the activation loop position in structures of apo-p38-2p (Fig. 2C), similar to a previous structure of BIRB796 bound to unphosphorylated p38α that showed a fully disordered activation loop^38^ (Fig. S6; PDB: 1KV2). Correlated with reduced WIP1 stimulation, the flipped activation loop conformation appears less stable as indicated by lack of coordination of the phosphate of pY182 by H174 and T175, which are not resolved in the electron density maps of the BIRB796-p38α-2p structure (Fig. S5E). Because BIRB796 and pexmetinib only differ in the portion of the molecule that docks in the ATP binding site of p38α (Fig. 3B & S2), we speculate that differences in this moiety cause two changes that account for differential stabilization of the flipped activation loop. First, the phenyl-ring of F169 interacts with the indazole of pexmetinib in contrast to a non-aromatic carbon of BIRB796. Second, the compounds interact differently with the linker (107-113) connecting the N and C-lobes of the kinase (Fig. 3A & E). This linker and the αD-helix that follows is ordered and resolved in the structure of p38α-2p bound to pexmetinib, with contacts between the terminal-hydroxyl group of pexmetinib with M109 and A111. In contrast, this region (111-118) is poorly ordered or unresolved for BIRB796-p38α-2p (Fig. 3E). Additionally, the αD-helix is unwound and there is a disulfide bond between C119 and C162^39^ (Fig. S7). Together, our results reveal that activation loop flipping promotes dephosphorylation of p38α-2p by WIP1 and suggest that the magnitude of phosphatase stimulation correlates with the stability of the flipped activation loop conformation. Further, our structures inform a model for how chemical groups of the inhibitors determine the magnitude of WIP1 stimulation.

**Figure 3:**
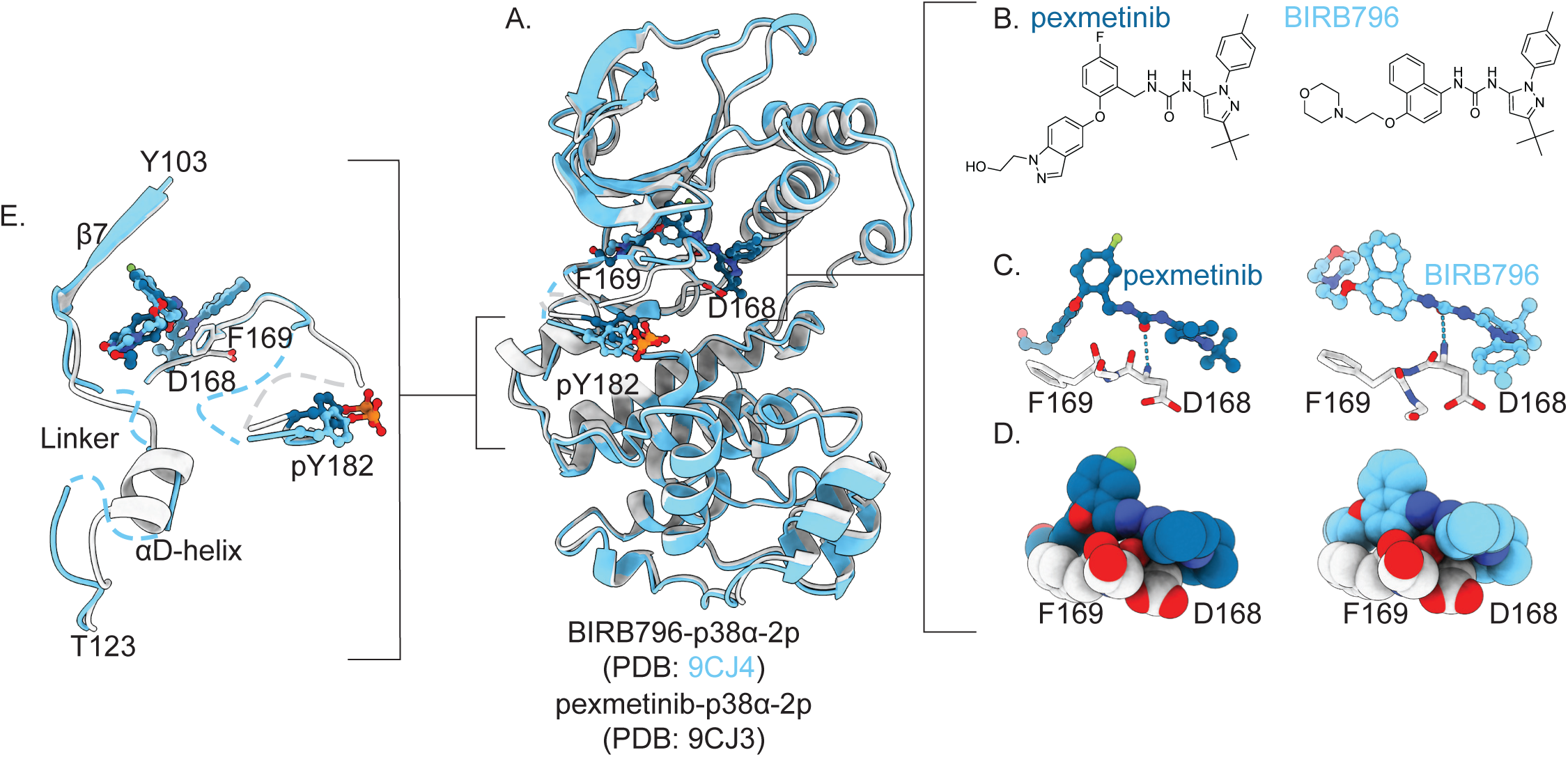
pY182 coordination in BIRB796-p38α-2p is structurally similar yet less stable than in pexmetinib-p38α-2p. (A) Overlay of BIRB796-p38α-2p (light blue; PDB: 9CJ4) and pexmetinib-p38α-2p (white, pexmetinib in dark blue; PDB: 9CJ3). D168 and F169 are shown as sticks and pY182 is shown as spheres and sticks. (B) Chemical structures of pexmetinib and BIRB796. (C) Interactions of pexmetinib (left; white and dark blue) and BIRB796 (right; white and light blue) with D168 and F169 (DFG motif) are shown in sticks with the hydrogen bond of the ligand to D168 shown as light blue dashes and (D) as spheres representing stacking and van der Waals interactions. (E) Zoom of linker and αD-helix region of BIRB796-p38α-2p (light blue) and pexmetinib-p38α-2p (white and dark blue). Dashed lines indicate regions that lacked electron density indicating increased flexibility for the BIRB796-p38α-2p structure. In all structures, oxygen, nitrogen, phosphorous and fluorine atoms are colored red, blue, orange and light green, respectively. Crystal contacts are illustrated in Fig. S13.

### A shared conformational state directs p38α dephosphorylation

To determine if any of the hundreds of structures of p38 in the PDB share the flipped activation loop conformation of pexmetinib and BIRB796-bound p38α-2p, we performed a DALI search^40^. This revealed a structure of human p38α-2p bound to DP802, a compound that lacks an ATP-binding site element but has a similar moiety that projects into the activation loop docking site as do pexmetinib and BIRB796 (Fig. S8A-B; PDB: 3NNX, RMSD 0.4 Å to pexmetinib-p38α-2p)^41^. Similar to the case of BIRB796 bound to p38α-2p, H174 and T175 do not coordinate the phosphate of pY182 and the αD-helix is disordered (Fig. S8A & C). Although DP802 was not commercially available for phosphatase assays, the authors noted that p38α-2p phosphorylation decreased in human cells treated with DP802 indicating a potential increase in cellular phosphatase activity^41^. Together with the structure, this finding supports our hypothesis that the shared moieties of pexmetinib, BIRB796, and DP802 are sufficient for activation loop flipping and predicts that DP802 would stimulate p38α dephosphorylation to some extent.

A search of deposited kinase structures using KinCoRe^42^ revealed one additional structure with a similar activation loop conformation: unphosphorylated human p38β bound to the ABL inhibitor nilotinib (Tasigna, PDB:3GP0, p38β shares 73.6% sequence identity to p38α). Based on this observation, we solved a structure of nilotinib-p38α-2p. The activation loop was indeed flipped similar to what we observed with our two dual-action inhibitors (S5C & F & S9A & B, Table S1; PDB: 9CJ1). Consistent with our prediction that the flipped activation loop presents the phospho-threonine for dephosphorylation, nilotinib stimulated WIP1 dephosphorylation of p38α-2p 5-fold (Fig. 1B-D & S1A & B). Thus, we conclude that nilotinib is a third dual-action inhibitor of p38α and that the resulting activation loop flipped conformation favors dephosphorylation by WIP1.

Although nilotinib originated from an unrelated compound series to that of pexmetinib and BIRB796, the chemical features that flip the activation loop are similar (Fig. S9B & S2). D168 is coordinated by a carbonyl oxygen of nilotinib and F169 makes hydrophobic interactions with nilotinib to anchor the N-terminus of the activation loop. Nilotinib has a moiety that displaces the activation loop, projecting even further beyond the ATP-binding pocket than pexmetinib and BIRB796 (Fig. S9B). Interestingly, imatinib (Gleevec), which does not stimulate WIP1 dephosphorylation of p38α-2p only differs from nilotinib in the moiety that projects beyond the ATP binding pocket (Fig. 1B, S1A & S2), emphasizing the importance of this moiety for phosphatase stimulation.

### Phospho-tyrosine is not required to stimulate threonine dephosphorylation

The fact that nilotinib causes activation loop flipping of unphosphorylated p38β (Fig. S9A) and that the coordination of the phosphate of pY182 is variable in our three structures of inhibitor-bound p38α-2p (Fig. S9C-E) raises the question of whether tyrosine phosphorylation is required for dual-action inhibition of p38α. We therefore tested whether our three dual-action compounds stimulate WIP1 dephosphorylation of singly threonine phosphorylated p38α^Y182F^-pT (Fig. 4A-B & S1A). Suggestive of a contribution of pY182 to dephosphorylation, WIP1 dephosphorylation of unliganded p38α^Y182F^-pT was 2-fold slower than for p38α-2p. While the stimulatory effects of BIRB796 and nilotinib on WIP1 dephosphorylation of p38α^Y182F^-pT were unchanged, the effect of pexmetinib was reduced an additional 2-fold (Fig. 4A-B & S1A). This result is consistent with the inference that additional contacts from tyrosine phosphorylation stabilize the flipped activation loop conformation of p38α when bound to pexmetinib and that such stabilization leads to an increased dephosphorylation rate.

**Figure 4:**
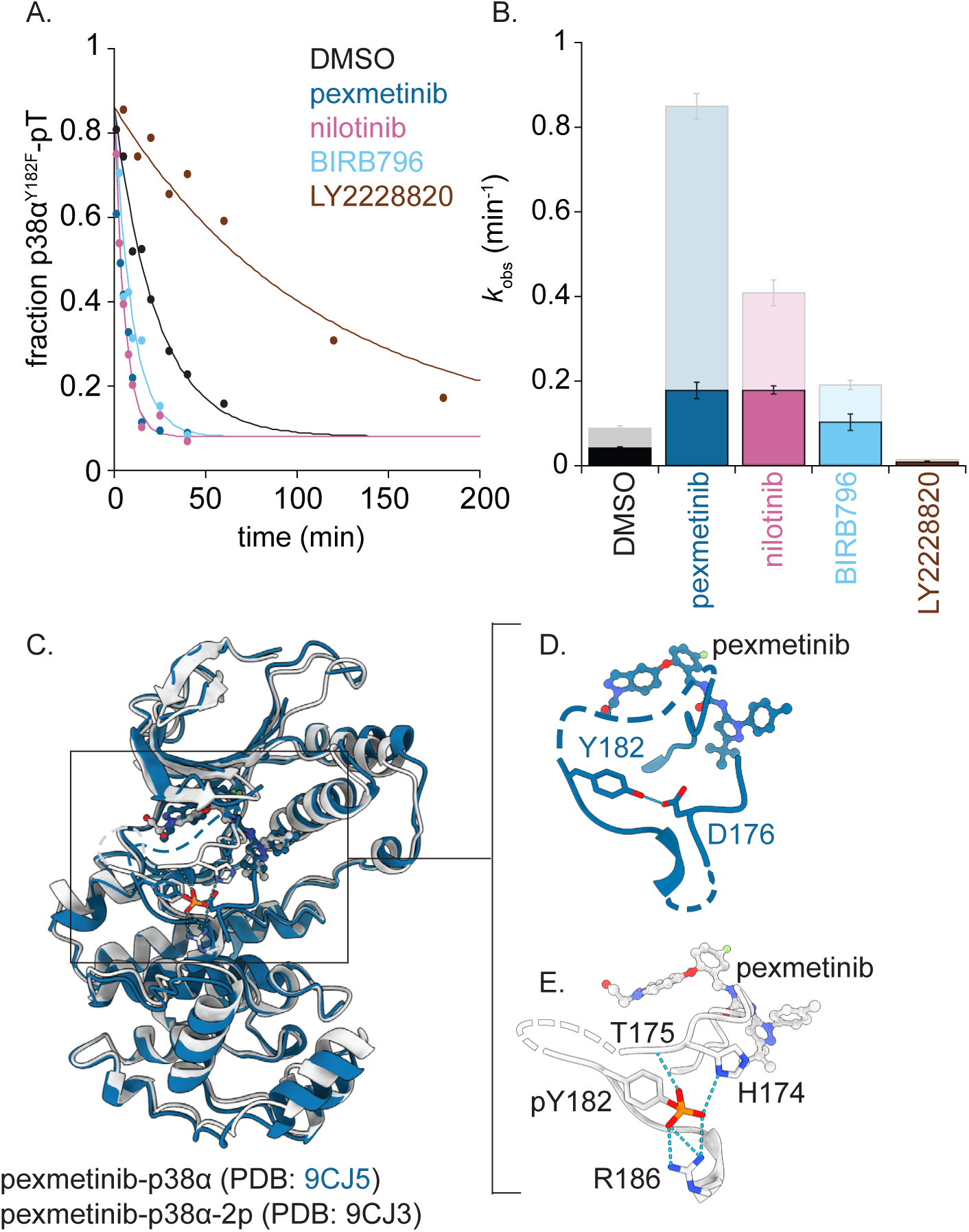
Tyrosine 182 phosphorylation is not required for stimulation of WIP1. (A) Dephosphorylation of p38α^Y182F^-pT by WIP1 performed under single turnover conditions: 2.5 µM WIP1 with 0.25 µM single threonine phosphorylated p38α^Y182F^-pT in the absence or presence of excess p38α inhibitors (1.25 µM). Data were fit to an exponential decay (see methods section, n=2 ± error of the fit). (B) Bar graph of rates derived from panel A (n=2 ± error of the fit) with *k*_obs_ of p38α-2p pT180 dephosphorylation by WIP1 (Fig. 1D) shown as translucent bars for reference. (C) X-ray crystal structure of unphosphorylated p38α bound to pexmetinib (dark blue; PDB: 9CJ5) overlayed with p38α-2p bound to pexmetinib (white; PDB: 9CJ3), showing a similar activation loop conformation. (D) Zoom in of pexmetinib-p38α activation loop (white) showing coordination of Y182. (E) Zoom in of pexmetinib-p38α-2p activation loop (dark blue) showing coordination of pY182. In all structures, blue dashes indicate hydrogen bonding and oxygen, nitrogen, phosphorous and fluorine atoms are colored red, blue, orange and light green, respectively. Crystal contacts are illustrated in Fig. S13.

Further demonstrating that tyrosine phosphorylation stabilizes activation loop flipping by pexmetinib, an X-ray crystal structure of unphosphorylated pexmetinib-p38α revealed that the unphosphorylated activation loop is flipped, with Y182 similarly positioned as pY182 (Fig. 4C-E & Table S1; PDB: 9CJ5). However, the resolution of the pexmetinib-p38α structure was significantly lower than for pexmetinib-p38α-2p, indicative of increased conformational flexibility. Together, we conclude that tyrosine phosphorylation promotes, but is not required for, pexmetinib-induced presentation of the p38α phospho-threonine for dephosphorylation by WIP1.

### Conformationally selective activation loop recognition is shared across phosphatase families

Since the conformational change of the activation loop that stimulates WIP1 dephosphorylation exposes pT180, we reasoned that other phosphatases that share a requirement for pT180 accessibility^34,37,43–45^ (Fig. S10), might similarly be stimulated by the dual-action compounds. We surveyed the subset of inhibitors that had the largest effects with three additional phosphatases: PPM1A, a PPM phosphatase related to WIP1 that natively targets p38α^25^, DUSP3, a dual specificity family phosphatase (related to tyrosine phosphatases that targets Ser/Thr and Tyr) that is capable of dephosphorylating p38α and has been well characterized biochemically^46^, as well as the alkaline phosphatase from shrimp (SAP). The same trends of dephosphorylation stimulation and inhibition were observed for all of the phosphatases but not to the same extent as seen for WIP1 (Fig. 5A, S1A & S11A-C). SAP had the largest change in activity upon inhibitor binding, followed by PPM1A and DUSP3. To determine if this effect was specific for threonine dephosphorylation, we assayed DUSP3 dephosphorylation of singly tyrosine phosphorylated p38α^T180A^-pY in the presence of our five modulating compounds. While the compounds induced small changes in dephosphorylation, the direction and magnitude of the effects were distinct from that for pT180 dephosphorylation of both p38α-2p and p38α^Y182F^-pT (Fig. 5B & S11D). Thus, the modulatory effects of conformation-selective inhibitors on p38α dephosphorylation are generalizable across phosphatase families, but the magnitude of change is specific to particular phosphatase/kinase pairs.

**Figure 5:**
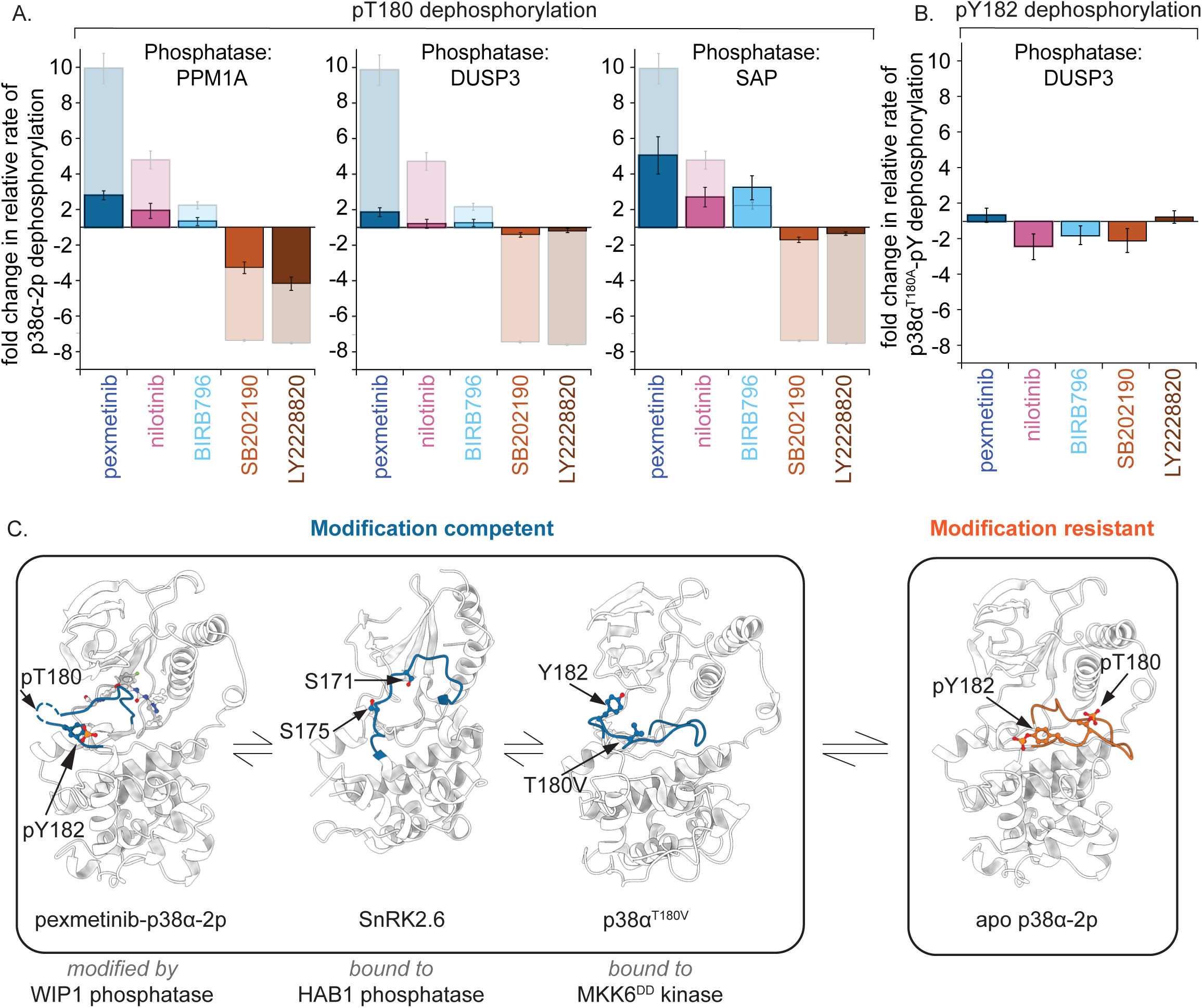
p38α exists in a conformational equilibrium that facilitates dephosphorylation by multiple phosphatases. (A) Comparisons of p38α-2p (0.25 µM) dephosphorylation rates by PPM1A (0.5 µM), DUSP3 (50 µM), and SAP (1 µM) in the presence of excess inhibitors (1.25 µM) as fold change compared to a DMSO control (error displayed is the propagated error of the fit). Comparable relative rates of p38α-2p dephosphorylation by WIP1 (Fig. 1B) are shown as translucent background bars on each plot as a reference. (B) Comparisons of pY182 of p38α^T180A^-pY (0.25 µM) dephosphorylation rates by DUSP3 (15 µM) in the presence of excess inhibitor (1.25 µM) compared to a DMSO control (error displayed is the propagated error of the fit (n=2)). (C) Model proposing a conformational equilibrium between modification resistant (human apo p38α-2p; PDB: 9CJ2) with inaccessible phosphorylation states, and modification competent states that represent exposed phosphorylation sites (pexmetinib-p38α-2p (PDB: 9CJ3), HAB1-SnRK2.6, HAB1 not shown (PDB: 3UGJ), and MKK6^DD^-p38α^T180V^, MKK6^DD^ not shown (PDB: 8A8M)).

## Discussion

Here, we have identified dual-action inhibitors of p38α MAP kinase that simultaneously inhibit the kinase by binding to the active-site and promote inactivation through dephosphorylation of the activation loop. This dual-action is accomplished by flipping the activation loop to a new conformation that presents the phospho-threonine for dephosphorylation. These discoveries provide answers to two longstanding questions: (1) how do phosphatases dephosphorylate their substrates when the phosphates are shielded from nucleophilic attack? and (2) how can cellular phosphatases be leveraged to enhance the efficacy of kinase inhibition?

### Phosphatases recognize a modification competent state of p38α-2p

How can active p38α-2p be turned off by phosphatases, given that the phosphorylated residues of the activation loop are inaccessible for nucleophilic attack as observed in X-ray crystal structures (Fig. S4 & S10)? Our new structures of p38α-2p bound to dual-action phosphatase-stimulating inhibitors delivered an answer to this puzzle: namely, that these inhibitors shift a pre-existing equilibrium of the activation loop^47^ towards a higher population of a dephosphorylation-competent state (Fig. 5C). Our biochemical and structural data suggest that this state exposes pT180 for dephosphorylation in an activation loop flipped conformation (Fig. 2A). While there are no structures yet of p38α bound to a cognate phosphatase, there exists a structure of unphosphorylated SnRK2.6 kinase bound to its cognate phosphatase, HAB1 (PDB: 3UJG, a PPM phosphatase related to WIP1)^48^. In this complex the activation loop is oriented to project outwards from the kinase in a related orientation to the flipped activation loop of p38α-2p bound to dual-action inhibitors, providing a template of how activation loop flipping could position the target residue for a docked phosphatase (Fig. 5C & S12A).

Suggestively, a recent cryo-EM structure of the upstream kinase (MKK6^DD^) trapped in complex with the substrate analogue p38α^T180V^ (PDB: 8A8M), similarly showed the p38α^T180V^ activation loop redirected towards the orientation that we report for p38α-2p bound to dual-action inhibitors^49^ (Fig. 5C & S12B). Whereas the activation loop conformation in our human apo p38α-2p structures and the published murine p38α-2p structure^33^ are resistant to dephosphorylation, binding of pexmetinib, BIRB796, or nilotinib shifts the equilibrium towards states that are dephosphorylation-competent (Fig. 5C). These dephosphorylation-competent states are similar, yet not identical, to the aforementioned structures of p38α^T180V^ and SnRK2.6 bound to their respective partners. This leads us to postulate that the activation loop of p38α samples an ensemble of related, but distinct, modification-competent states that determine the extent to which p38α activation loop is phosphorylated and dephosphorylated (Fig. 5C). Because conformational flexibility of the activation loop is a general feature of protein kinases^3^, it is possible that similar conformational equilibria determine how protein phosphatases target diverse kinases.

### Dual-action kinase inhibitors may increase drug efficacy by leveraging phosphatases

How can cellular phosphatases be leveraged to enhance the efficacy of kinase inhibition? A dual-action kinase inhibitor such as pexmetinib, nilotinib, or BIRB796 that stimulates dephosphorylation of its target could achieve increased efficacy. This could be achieved by increasing the potency or completeness of inhibition, or by blocking aspects of the kinase mechanism that are phosphorylation dependent but do not require kinase activity. In contrast to previous approaches that used heterobifunctional molecules ^10,11,13^, our discovery reveals that both modes of action can be achieved by a single compound that traps a phosphatase preferred conformational state. This provides a new roadmap to identifying dual-action kinase inhibitors. Finally, cell-type variability of phosphatase activity profiles raises the possibility that specificity conferred by such dual-action inhibition mechanism could enable rationally guided identification of cell-type specific kinase inhibitors. Suggestive that similar dual-action inhibitors may be identifiable for diverse kinases, inhibitors that trap the activation loop of the MAP kinase ERK2 in an inactive conformation (distinct from the flipped-p38α conformation we observe) stimulate ERK tyrosine dephosphorylation^50^, and some Akt inhibitors block dephosphorylation by trapping an active state of Akt that occludes the phosphorylated residues^51^.

Our finding that nilotinib and pexmetinib stimulate p38α dephosphorylation may provide examples of the clinical relevance of dual-action inhibition. Initial biochemical kinase screening identified p38α as a weaker off-target binder of nilotinib (*K*_d_: 460 nM) compared to its primary target ABL (*K*_d_: 10 nM)^52^. However, nilotinib’s side effects correlated with p38 inhibition^53^, which could be exacerbated by its phosphatase interaction. Conversely, nilotinib has been investigated as a treatment for neurodegenerative diseases including Lewy Body Dementias^54^, Alzheimer’s disease^55,56^, and Parkinson’s disease^57^, with therapeutic benefit attributed to reduction of inflammation in the brain that could be due to p38 inhibition^58^. Similarly, pexmetinib is currently in a phase II clinical trial for solid tumors and it is possible that interactions with phosphatases in this context could impact clinical outcome. Thus, our discovery that conformation-selective kinase inhibitors can control phosphatase activity towards their targets could enable the development of therapeutics with improved efficacy and emphasizes the importance of determining how kinase conformations control their dephosphorylation.

## Acknowledgements

The authors thank Chris Miller, Julia Kardon, Richard Losick, Liz Hedstrom, Rachelle Gaudet, Mike Marr, Fred Raab III, and members of the Bradshaw Lab for input, inspiration, and/or critical reading of the manuscript. We additionally benefited from input from the SPROUT program and judges organized by the Brandeis Office of Technology Licensing and the NSF I-Corps program. N.B. received startup funds from Brandeis University, support from the Brandeis SPROUT program, and support from the NSF I-Corps Site 1644666NSF. D.K. is supported by the Howard Hughes Medical Institute (HHMI). The Berkeley Center for Structural Biology is supported by HHMI, Participating Research Team members, and the NIH, National Institute of General Medical Sciences, ALS-ENABLE grant P30 GM124169. The Advanced Light Source is a Department of Energy Office of Science User Facility under Contract No. DE-AC02-05CH11231. The Pilatus detector on beamline 2.0.1 was funded under NIH grant S10OD021832.

## Author contributions

E.S. and N.B. designed research; E.S., H.L., R.K., and X.W. performed research; E.S., X.W., and Y.Q. contributed new reagents/analytic tools; E.S., H.L., R.K., X.W., and N.B. analyzed data, E.S., H.L., D.K., and N.B. wrote the paper.

## Methods

### Protein expression constructs

Full length PPM1A, p38α, and DUSP3 were generated by isolation of their respective coding sequences from HEK 293 genomic DNA and inserted into pET47b vectors with an N-terminal 6-His tag. Mutations were introduced to the p38α expression construct using the QuikChange site-directed mutagenesis kit (Agilent). pCDFDuet-MKK6-EE was a gift from Kevin Janes (Addgene plasmid #82718; http://n2t.net/addgene:82718; RRID: Addgene 82718). The cloning sequence was inserted into a pET47b vector with an N-terminal 6-His tag. The 1-420 WIP1 construct was codon optimized by GenScript and was inserted into a pET28b vector with an N-terminal 6-His SUMO tag.

### Protein expression and purification

All proteins were expressed in *E. coli* BL21 (DE3) cells, were grown at 37 °C in Lennox lysogeny broth (LB) to an OD600 of 0.6 and induced at 16 °C for 14-18 hours with 1 mM isopropyl β-d-1-thiogalactopyranoside (IPTG) unless otherwise specified. Cells were harvested and purified as followed:

#### WIP1

In addition to 1 mM IPTG cells were induced with 2 mM MgCl_2_. Cell pellets were resuspended in lysis buffer (50 mM Tris-HCl pH 7.4, 500 mM NaCl, 10 mM MgCl_2_, 10% (v/v) glycerol, 1 mM dithiothreitol (DTT)) with 1 mM phenylmethylsulphonyl fluoride (PMSF), 10 μg/mL lysozyme and 1:1000 (by volume) benzonase and were lysed using three passes in a microfluidizer at 10,000 PSI. Cell lysates were cleared by spinning at 16,000 RPM for 45 minutes in an Avanti JA-20 rotor. 10 mM imidazole was added to the cleared lysates. A HisTrap HP column on an AKTA FPLC was equilibrated with lysis buffer and 6% elution buffer (50 mM Tris-HCl pH 7.4, 500 mM NaCl, 10 mM MgCl_2_, 10% (v/v) glycerol, 1 mM DTT, 400 mM imidazole). Cleared lysates were then run over a HisTrap HP column, washed with 6% elution buffer for 10 column volumes, and eluted over a gradient to 100% elution buffer over 20 column volumes. Purity of fractions was analyzed via SDS-PAGE using a 10% Tris-Tricine polyacrylamide gel stained with Coomassie brilliant blue solution. Protein containing fractions were pooled and the SUMO-His tags were cleaved with 10 μg per mg total protein of ULP1-R3 protease^59^ in dialysis to lysis buffer overnight at 4 °C. WIP1 was further purified on a Superdex S200 16/600 column equilibrated with lysis buffer. Fractions were pooled, concentrated to 200 μM and treated with a 5-fold molar excess of EDTA to remove metal. Chelated WIP1 was buffer exchanged into storage buffer (50 mM Tris-HCl pH 7.4, 500 mM NaCl, 10% glycerol (v/v)), flash-frozen and stored at -80 °C.

#### PPM1A, MKK6^EE^, p38α and p38α mutants

Cell pellets were resuspended in lysis buffer (50 mM HEPES pH 7.5, 200 mM NaCl, 20 mM imidazole, 10% (v/v) glycerol, 0.5 mM DTT) with 1 mM PMSF and were lysed using three passes in a microfluidizer at 10,000 PSI. Cell lysates were cleared by spinning at 16,000 RPM for 45 minutes in an Avanti JA-20 rotor. A HisTrap HP column on an AKTA FPLC was equilibrated with lysis buffer. Cleared lysates were then run over a HisTrap HP column, washed with lysis buffer for 10 column volumes, and eluted over a gradient to 100% elution buffer elution buffer (50 mM HEPES pH 7.5, 200 mM NaCl, 400 mM imidazole, 10% (v/v) glycerol, 0.5 mM DTT)) over 20 column volumes. Purity of fractions was analyzed via SDS-PAGE using a 10% Tris-Tricine polyacrylamide gel stained with Coomassie brilliant blue solution. Protein containing fractions were pooled and the 6-His tags were cleaved with 3C protease at a 1:100 protease to protein molar ratio in dialysis to lysis buffer overnight at 4 °C. Cleaved tags were subtracted by passing over a column containing Ni-NTA resin equilibrated with lysis buffer. Proteins were further purified on a Superdex S200 16/600 column equilibrated with FPLC buffer (50 mM HEPES pH 7.5, 200 mM NaCl, 10% (v/v) glycerol, 2 mM DTT). Fractions were pooled, concentrated to 500 μM, flash-frozen and stored at -80°C.

#### DUSP3

Cell pellets were resuspended in lysis buffer (50 mM HEPES pH 7.5, 200mM NaCl, 20 mM imidazole, 10% (v/v) glycerol, 0.5 mM DTT) with 1 mM PMSF and were lysed using three passes in a microfluidizer at 10,000 PSI. Cell lysates were cleared by spinning at 16,000 RPM for 45 minutes in an Avanti JA-20 rotor. A HisTrap HP column on an AKTA FPLC was equilibrated with lysis buffer. Cleared lysates were then run over a HisTrap HP column, washed with lysis buffer for 10 column volumes, and eluted over a gradient to 100% elution buffer elution buffer (50 mM HEPES pH 7.5, 200 mM NaCl, 400 mM imidazole, 10% (v/v) glycerol, 0.5 mM DTT) over 20 column volumes. Fractions containing protein were pooled, concentrated to 800 μM, flash-frozen and stored at -80°C.

### Small molecule reagents

| Compound | Source | Cat. # |
| --- | --- | --- |
| pexmetinib | APExBIO (Houston, TX) | B6012 |
| nilotinib | APExBIO (Houston, TX) | A8232 |
| BIRB796 | APExBIO (Houston, TX) | A5639 |
| imatinib | APExBIO (Houston, TX) | A3487 |
| SD169 | APExBIO (Houston, TX) | C5850 |
| VX-702 | APExBIO (Houston, TX) | A8687 |
| FGA-19 | Fisher Scientific (Hampton, NH) | 5304860001 |
| SB203580 | APExBIO (Houston, TX) | A8254 |
| RWJ67657 | APExBIO (Houston, TX) | C5316 |
| VX-745 | APExBIO (Houston, TX) | A8686 |
| TAK-715 | APExBIO (Houston, TX) | A8688 |
| losmapimod | APExBIO (Houston, TX) | B4620 |
| SB202190 | APExBIO (Houston, TX) | A1632 |
| LY2228820 | APExBIO (Houston, TX) | A5566 |
| Fluorescein diphosphate, tetra ammonium salt (FDP) | Fisher Scientific (Hampton, NH) | 501953367 |

### X-ray crystallography

#### Crystallization

p38α was dual phosphorylated by MKK6^EE^ using established methods^47^. Dual phosphorylation was verified by the protein intact mass measurements using mass spectrometry.

Crystals of p38α-2p apo were obtained by combining 0.3 µL of 8 mg/mL p38α-2p with 0.3 µL of 100 mM BIS-TRIS pH 6.5 and 25% polyethylene glycol (PEG) 3350. Crystals were grown in sitting drops for two weeks at 20 °C. Crystals were harvested and flash-frozen (liquid N_2_) in 15% glycerol.

Crystals of p38α-2p bound to pexmetinib were obtained by combining 0.3 µL of 8 mg/mL p38α-2p + 250 µM pexmetinib for a final DMSO concentration of 5% with 0.3 µL of 100 mM MES pH 6.5, 200 mM ammonium sulfate, 4% 1,3-Propanediol, and 30% PEG8000. Crystals were grown in sitting drops for two weeks at 20 °C. Crystals were harvested and flash-frozen (liquid N_2_) in 15% glycerol.

Crystals of p38α-2p bound to nilotinib were obtained by combining 0.3 µL of 8 mg/mL p38α-2p + 250 µM nilotinib for a final DMSO concentration of 5% in a 0.3 µL reservoir of 100 mM BIS-TRIS pH 6.0 and 23% PEG3350. Crystals were grown in sitting drop for two months at 20 °C. Crystals were harvested and flash-frozen (liquid N_2_) in 15% glycerol.

Crystals of p38α-2p bound to BIRB796 were obtained by combining 0.3 µL of 8 mg/mL p38α- 2p + 250 µM BIRB796 for a final DMSO concentration of 5% with 0.3 µL reservoir of 100 mM BIS-TRIS pH 5.5, 200 mM ammonium sulfate, and 25% PEG3350. Crystals were grown in sitting drops for two weeks at 20°C. Crystals were harvested and flash-frozen (liquid N_2_) in 15% glycerol.

Crystals of p38α bound to pexmetinib were obtained by combining 0.3 µL of 8 mg/mL p38α + 250 µM pexmetinib for a final DMSO concentration of 5% in a 0.3 µL reservoir of 100 mM MES pH 6.0, 200 mM ammonium sulfate, 4% 1,3-Propanediol, and 20% PEG6000. Crystals were grown in sitting drops for two weeks at 20 °C. Crystals were harvested and flash-frozen (liquid N_2_) in 15% glycerol.

##### Data collection and processing

Cryogenic (100 K) X-ray diffraction data of single crystals were collected at Advanced Light Source (Lawrence Berkeley National Laboratory) at beamlines 2.0.1 (p38α-2p and pexmetinib- p38α-2p) and 8.2.2 (nilotinib-p38α-2p, BIRB796-p38α-2p and pexmetinib-p38α). The data were integrated with XDS^60^, scaled and merged in Aimless^61^, and data quality was assessed using Xtriage (Phenix)^62^. Structures of p38α-2p and pexmetinib-p38α-2p were solved by molecular replacement using Phaser (Phenix)^63^ utilizing as a search model chain A of 3PY3 (murine p38α- 2p). Initial phases for nilotinib-p38α-2p, BIRB796-p38α-2p and pexmetinib-p38α were obtained using pexmetinib-p38α-2p as a search model.

##### Refinement and model building

Refinement and manual model building were performed using phenix.refine (Version 1.20.1)^64^ and Coot,^65^ respectively. Models were validated using MolProbity^66^. Figures of structure models were created using ChimeraX^67^.

### Phosphatase assays

All phosphatase assays were performed with p38α-2p that was labeled with ^32^P by incubating p38α (25 µM), 6His-MKK6^EE^ (0.625 µM), and 20 µCi of γ-^32^P ATP for 6-8 hours at room temperature in 20 mM HEPES pH 7.5, 0.5 mM EDTA, 20 mM MgCl_2_, 2 mM DTT. Following initial incubation, excess cold ATP was added for a final concentration of 12 mM and was incubated overnight. This results in dual-phosphorylated p38α with the ^32^P_i_ nearly exclusively incorporated on T180. Unincorporated nucleotide was removed by buffer exchange using a Zeba spin column equilibrated in 50 mM HEPES pH 7.5, 100 mM NaCl. 6His-MKK6^EE^ was then removed by Ni-NTA resin equilibrated in 50 mM HEPES pH 7.5, 100 mM NaCl, 20 mM imidazole. The flow-through fraction from the Ni-NTA resin containing ^32^pT180 labeled p38α- 2p was then exchanged into 50 mM HEPES pH 7.5, 100 mM NaCl buffer using 3 subsequent Zeba spin columns to remove all unincorporated nucleotide and free phosphate. ^32^pT180 labeled p38α-2p was aliquoted and frozen at -80°C for future use.

All phosphatase assays were analyzed using the following exponential decay equation:

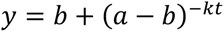

With a being the y-intercept, b being the baseline, k being the *k*_obs_, and t being time

#### WIP1 and PPM1A

All WIP1 phosphatase assays were performed at room temperature in 50 mM HEPES pH 7.5, 0.8 mM CHAPS, 0.05 mg/mL BSA, and 15 mM MnCl_2_. WIP1 concentration was 2.5 µM, PPM1A concentration was 0.5 µM and ^32^pT180 labeled p38α-2p was 0.25 µM unless otherwise stated. 1.25 μM p38α inhibitors were added to reactions at a 5% final DMSO concentration immediately before the start of the reaction. Reactions were stopped with 0.5 M EDTA pH 8.0 and run on PEI-Cellulose TLC plates developed in 1 M LiCl_2_ and 0.8 M acetic acid and imaged on a Typhoon scanner. Phosphatase assays were performed more than three independent times as separate experiments.

#### DUSP3

All DUSP3 phosphatase assays were performed in 50 mM HEPES pH 7.5 and 100 mM NaCl. Reactions were stopped with SDS and run on PEI-Cellulose TLC plates run through water then developed in 1 M LiCl_2_ and 0.8 M acetic acid and imaged on a Typhoon scanner. Phosphatase assays were performed more than three independent times as separate experiments. Data shown in figures is from a single representative experiment, and reported errors are the error from the fit unless indicated otherwise. Reactions with p38α-2p were run at 37 °C with 50 µM DUSP3 and 0.25 µM ^32^pT180 labeled p38α-2p. 1.25 µM p38α inhibitors were added to reactions at a 5% final DMSO concentration immediately before the start of the reaction. Reactions with ^32^pY182 labeled p38α^T180A^-pY were run at room temperature with 15 µM DUSP3 and 0.25 µΜ ^32^pY182 labeled p38α^T180A^-pY. 1.25 µM p38α inhibitors were added to reactions at a 5% final DMSO concentration immediately before the start of the reaction.

#### SAP

All SAP phosphatase assays were performed at 37 °C in 1x SAP reaction buffer. SAP concentration was 1 unit per 5 µL reaction and ^32^pT180 labeled p38α-2p concentration was 0.25 µM. 1.25 µM p38α inhibitors were added to reactions at a 5% final DMSO concentration immediately before the start of the reaction. Reactions were stopped with SDS and run on PEI-Cellulose TLC plates run through water then developed in 1 M LiCl_2_ and 0.8 M acetic acid and imaged on a Typhoon scanner. Phosphatase assays were performed more than three independent times as separate experiments. Data shown in figures is from a single representative experiment, and reported errors are the error from the fit unless indicated otherwise.

### Fluorescein diphosphate phosphatase assay

All FDP reactions were performed at 25 °C in a Corning 3573 384-well black flat bottom plate in 50 mM K HEPES pH 7.5, 0.8 mM CHAPS, 0.05 mg/mL BSA, 10 mM MnCl_2_, 30 nM WIP1 and 50 µM FDP. 1.25 µM p38α inhibitors were added at a final DMSO concentration of 5% immediately before the start of the reaction. Fluorescence (*λ*_ex_: 470 nm *λ*_em_: 530 nm) was read on a plate reader taking timepoints every 30 seconds at a constant temperature of 25 °C.

---

**Figure S1:**
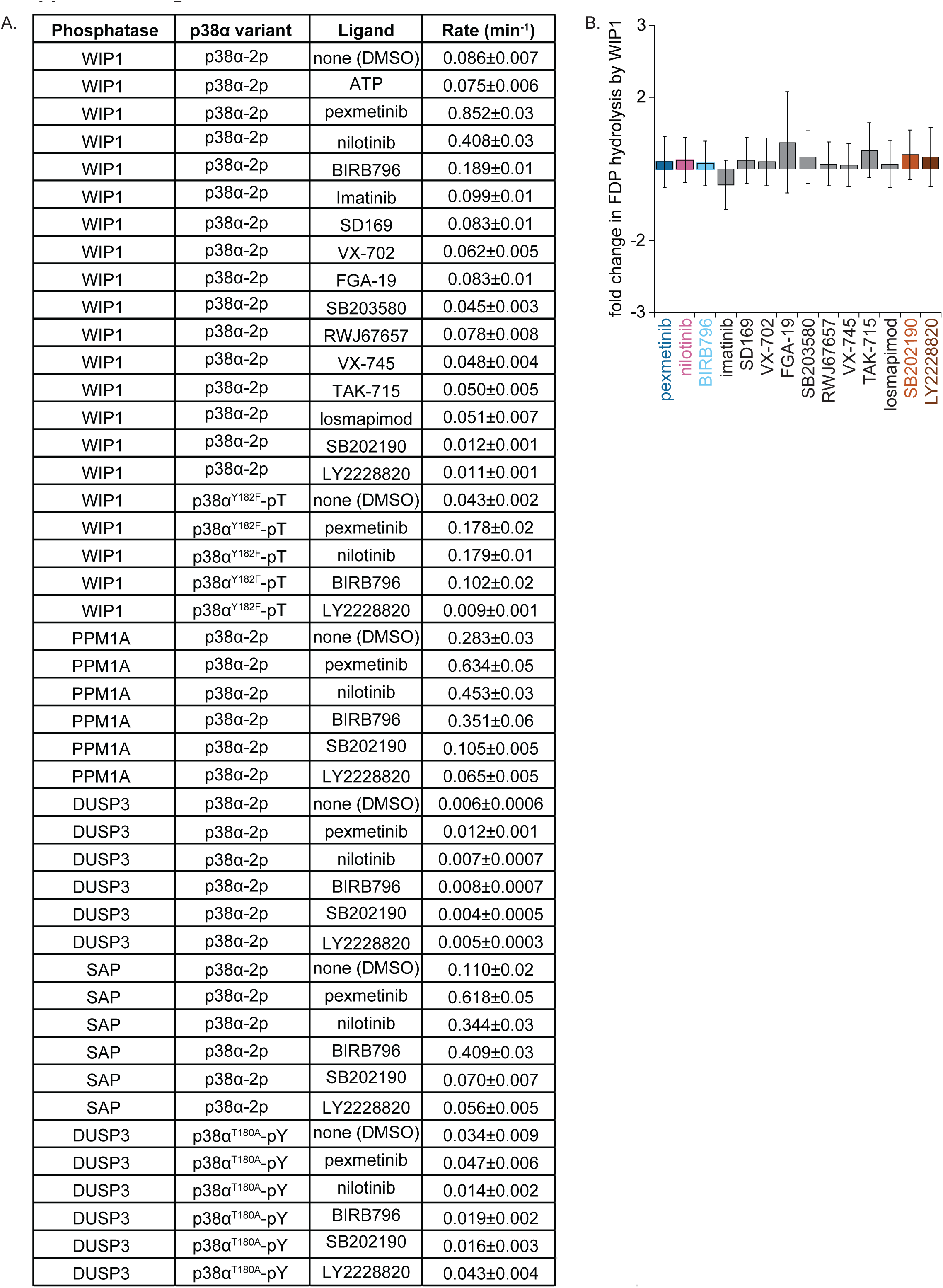
Single turnover kinetics compiled rates. (A) Table of compiled rates (*k*_obs_ (min^-1^)) ± error of the fit from data shown in Figures 1, 4, 5, and S11. (B) Multiple turnover kinetics of fluorescein diphosphate (FDP; 50 µM) hydrolysis by WIP1 (30 nM) in the absence and presence of 1.25 µM inhibitors showing no change in WIP1 activity toward a generic substrate with the addition of p38α inhibitors. Data is shown as the fold change in *k*_obs_ in the presence of inhibitors relative to a DMSO treated control (*k*_obs_ were averaged from an n=3 ± propagated 1 SD)

**Figure S2:**
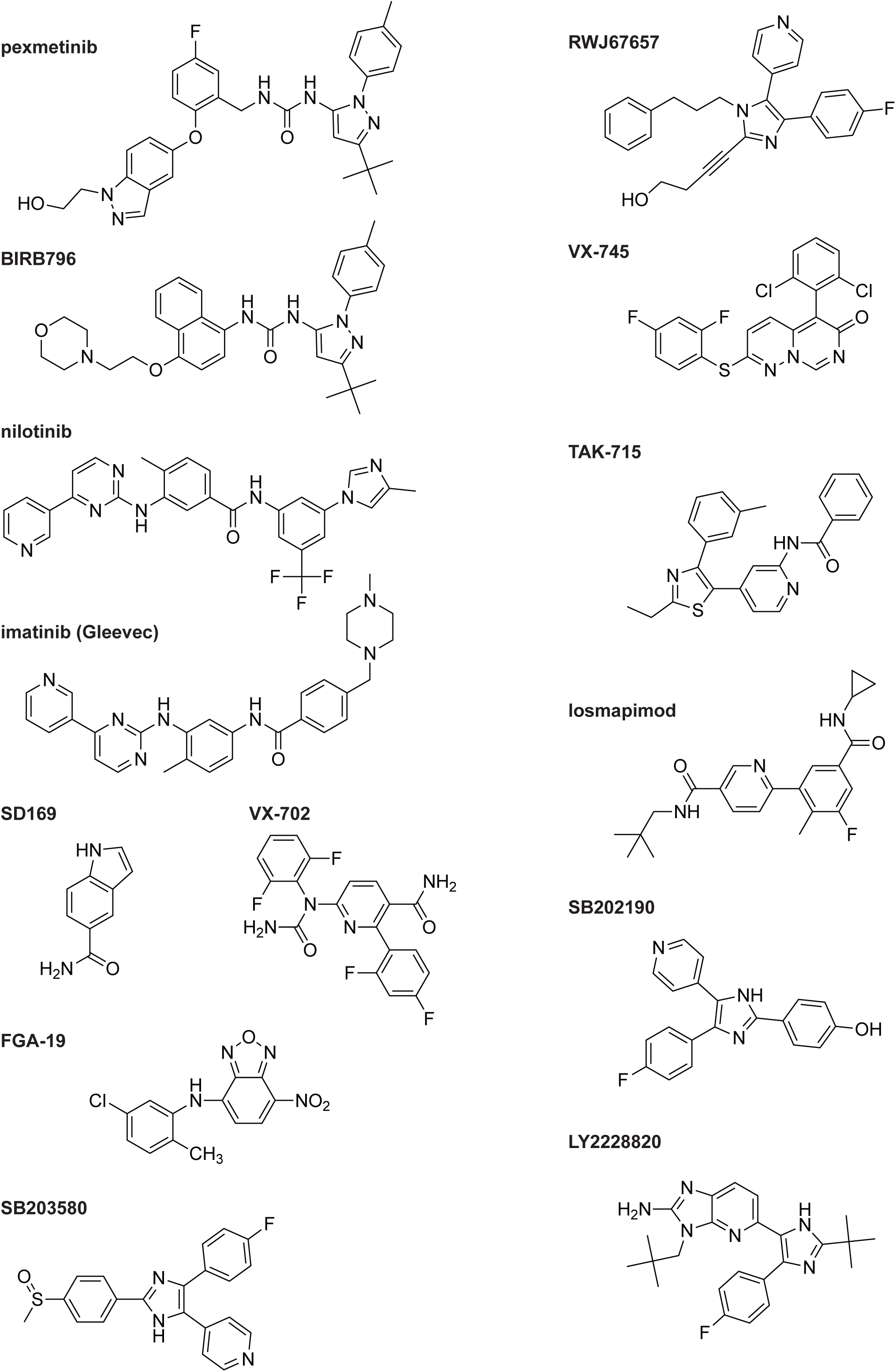
Chemical structures of all compounds. Chemical structures of compounds tested in Fig 1B.

**Figure S3:**
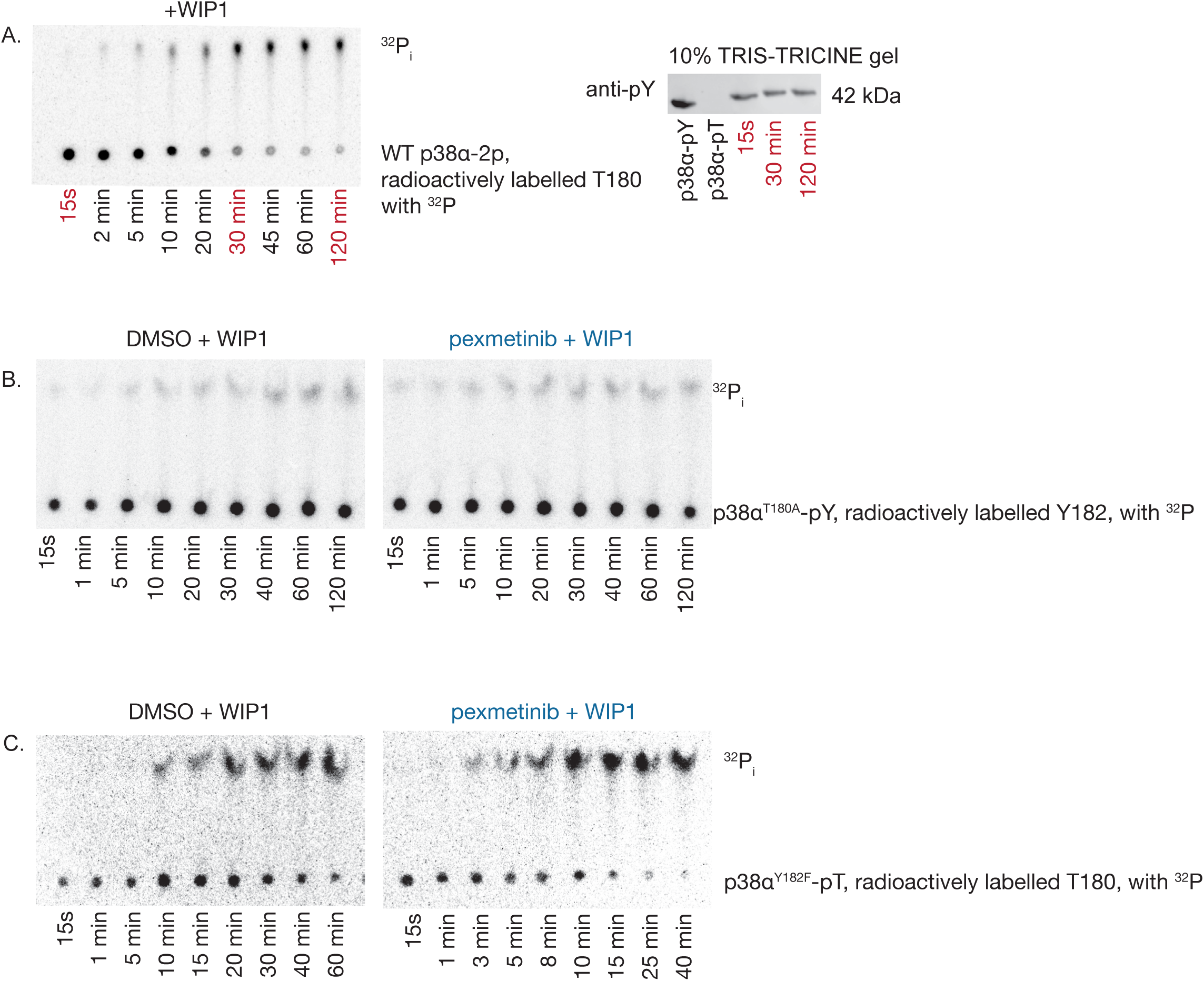
MKK6^EE^ preferentially phosphorylates T180 in p38α and WIP1 dephosphorylates only pT180 in p38a-2P. (A) Time course of WIP1 (2.5 µM) catalyzed p38α-2p (0.25 µM) dephosphorylation. Reaction is visualized on a TLC plate over time. p38α-2p (radioactively labelled pT180, with ^32^P) substrate is seen on the bottom and radioactive free phosphate (^32^P_i_) is the product at the top of the TLC plate (left panel). Time points in red are shown on a western blot on the right, probed for phospho-tyrosine, showing no phospho-tyrosine dephosphorylation over the reaction time course. This indicates that the sequential addition of sub saturating γ-^32^P ATP followed by excess cold ATP during the phosphorylation reaction by MKK6^EE^ results in a sample of p38α-2p which is radioactively labeled exclusively on pT180. Therefore, the measured rates in Fig. 1B-D of p38α-2p dephosphorylation are p38α-pT (radioactive pT180, with ^32^P) in the context of p38α-2p. (B) Dephosphorylation of p38α^T180A^-pY (0.25 µM, radioactive Y182, with ^32^P) by WIP1 (2.5 µM) in the absence (left) or presence (right) of excess pexmetinib (1.25 µM) shown on a TLC plate over time. p38α^T180A^-pY (with ^32^P) substrate is seen on the bottom and radioactive free phosphate is the product at the top of the TLC plate. WIP1 cannot dephosphorylate p38α^Y182F^-pT (with ^32^P). (C) Dephosphorylation of p38α^Y182F^-pT (0.25 µM with ^32^P) by WIP1 (2.5 µM) in the absence (left) or presence (right) of excess pexmetinib (1.25 µM) shown on a TLC plate over time. p38α^Y182F^-pT (with ^32^P) is seen on the bottom and radioactive free phosphate is the product at the top of the TLC plate.

**Figure S4:**
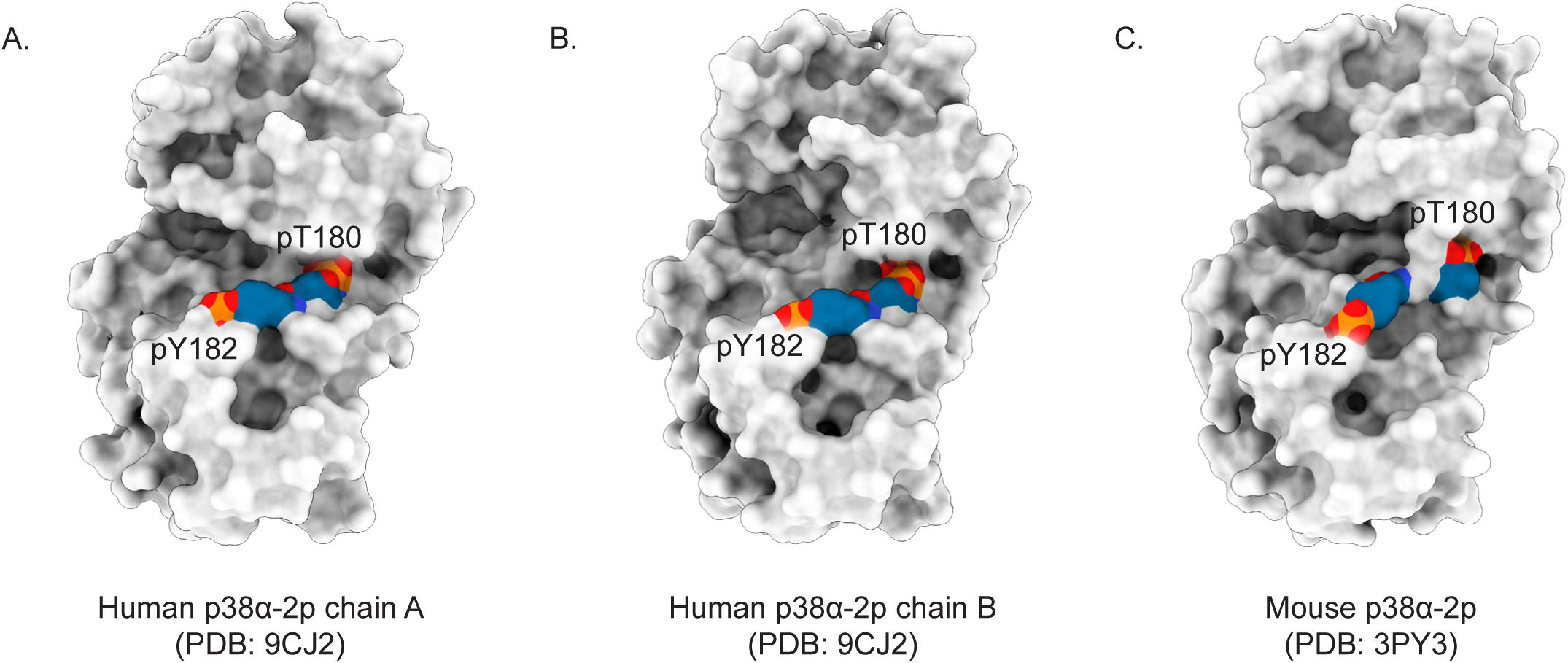
p38α-2p phosphorylation sites are inaccessible to nucleophilic attack. (A-C) Surface representation of human p38α-2p chain A (A) and chain B (B) (PDB: 9CJ2) and murine p38α-2p (PDB: 3PY3) showing buried phospho-sites inaccessible for nucleophilic attack by a phosphatase. In all structures, oxygen, nitrogen, phosphorous and fluorine atoms are colored red, blue, orange and light green, respectively. Crystal contacts are illustrated in Fig. S13.

**Figure S5:**
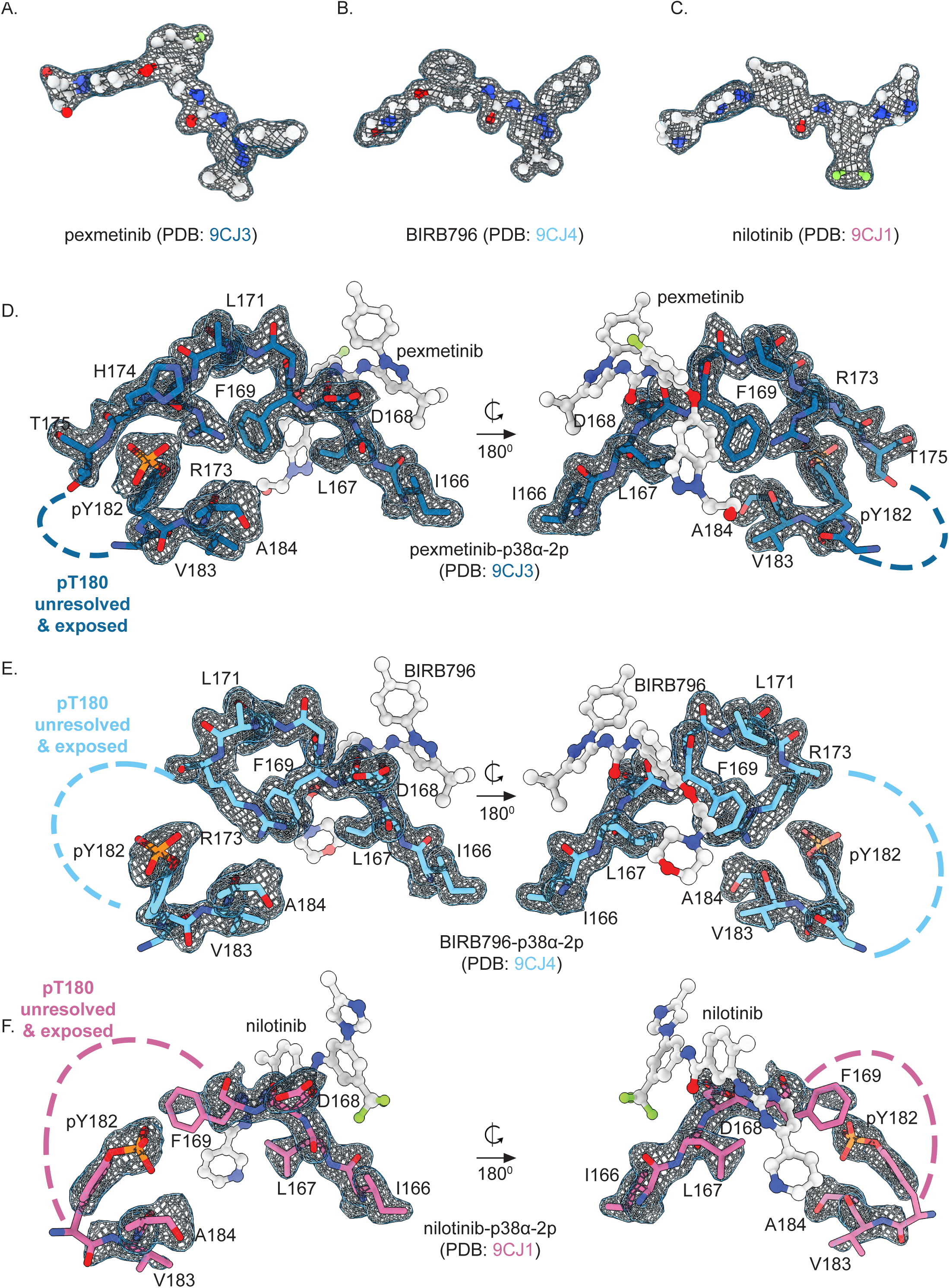
Activation loops of ligand bound p38α structures are resolved differently. 2*F*_o_-*F*_c_ electron density (grey mesh; contoured at 1 σ within 1.5 Å within selection) of (A) pexmetinib (PDB: 9CJ3) (B) BIRB796 (PDB: 9CJ4) and (C) nilotinib (PDB: 9CJ1). (D) 2*F*_o_-*F*_c_ electron density (grey mesh; contoured at 1 σ within 1.5 Å within selection) of activation loop (167-184) in p38α-2p bound to pexmetinib model (dark blue; PDB: 9CJ3). (E) 2*F*_o_-*F*_c_ electron density (grey mesh; contoured at 1 σ within 1.5 Å within selection) of activation loop (167-184) in p38α-2p bound to BIRB796 model (light blue; PDB: 9CJ4). (F) 2*F*_o_-*F*_c_ electron density (grey mesh; contoured at 1 σ within 1.5 Å within selection) of activation loop (167-184) in p38α-2p bound to nilotinib model (pink; PDB: 9CJ1). In all structures, oxygen, nitrogen, phosphorous and fluorine atoms are colored red, blue, orange and light green, respectively. Crystal contacts are illustrated in Fig. S13.

**Figure S6:**
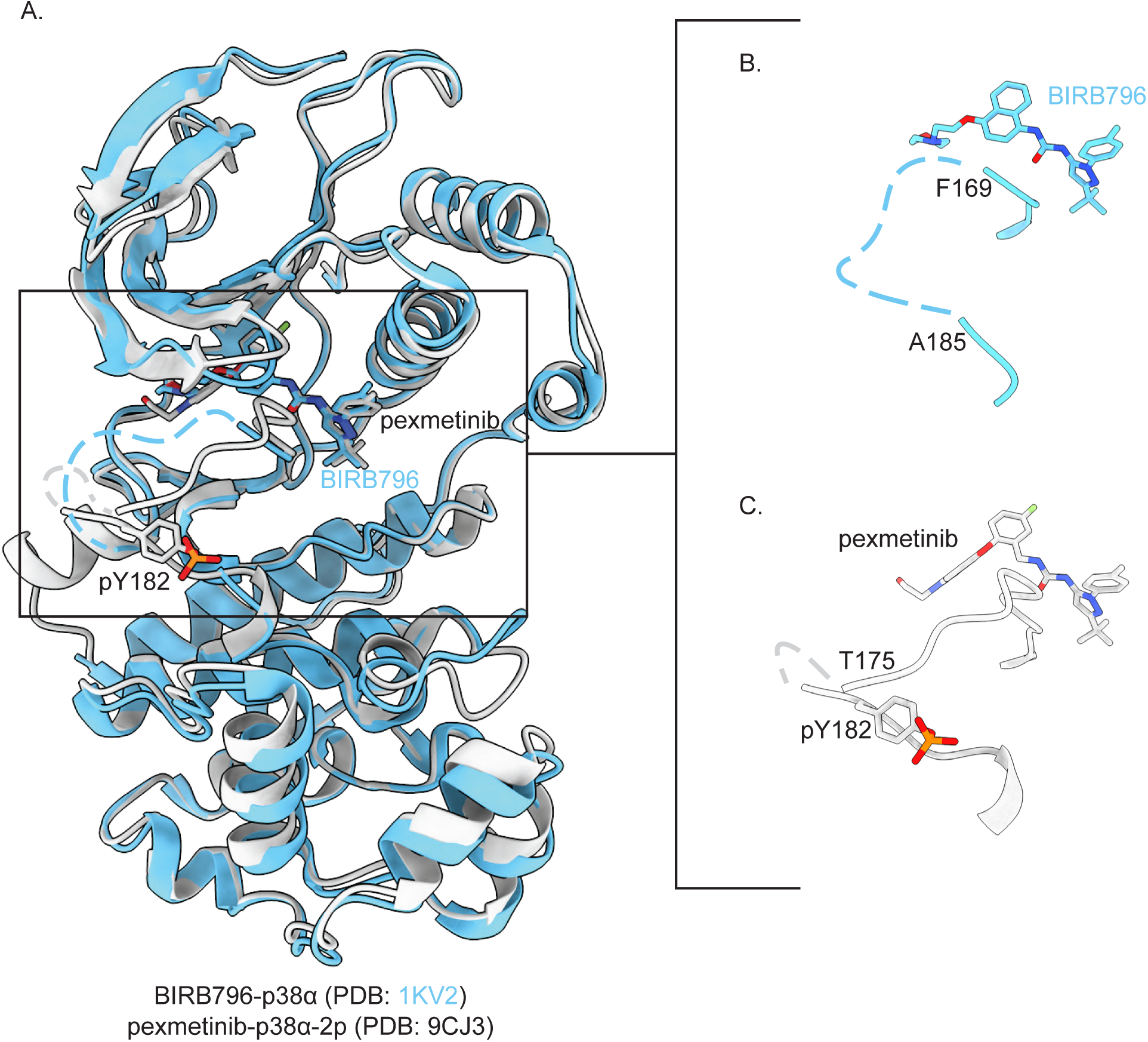
Pexmetinib binding to p38α-2p is similar to known binding of BIRB796 to unphosphorylated p38α. (A) Overlay of previously published crystal structure of unphosphorylated human p38α bound to BIRB796 (light blue; PDB: 1KV2) and our structure of dual phosphorylated human p38α bound to pexmetinib (white; PDB: 9CJ3) showing a similar binding mode between the two ligands, including missing electron density for BIRB796-p38α (residues 170-184) and pexmetinib-p38α-2p (residues 176- 180). Phospho-tyrosine in the pexmetinib-p38α-2p is shown as sticks. Zoom depicts the ligand binding site and activation loop of (B) BIRB796-p38α and (C) pexmetinib-p38α- 2p. In all structures, oxygen, nitrogen, phosphorous and fluorine atoms are colored red, blue, orange and light green, respectively. Crystal contacts are illustrated in Fig. S13.

**Figure S7:**
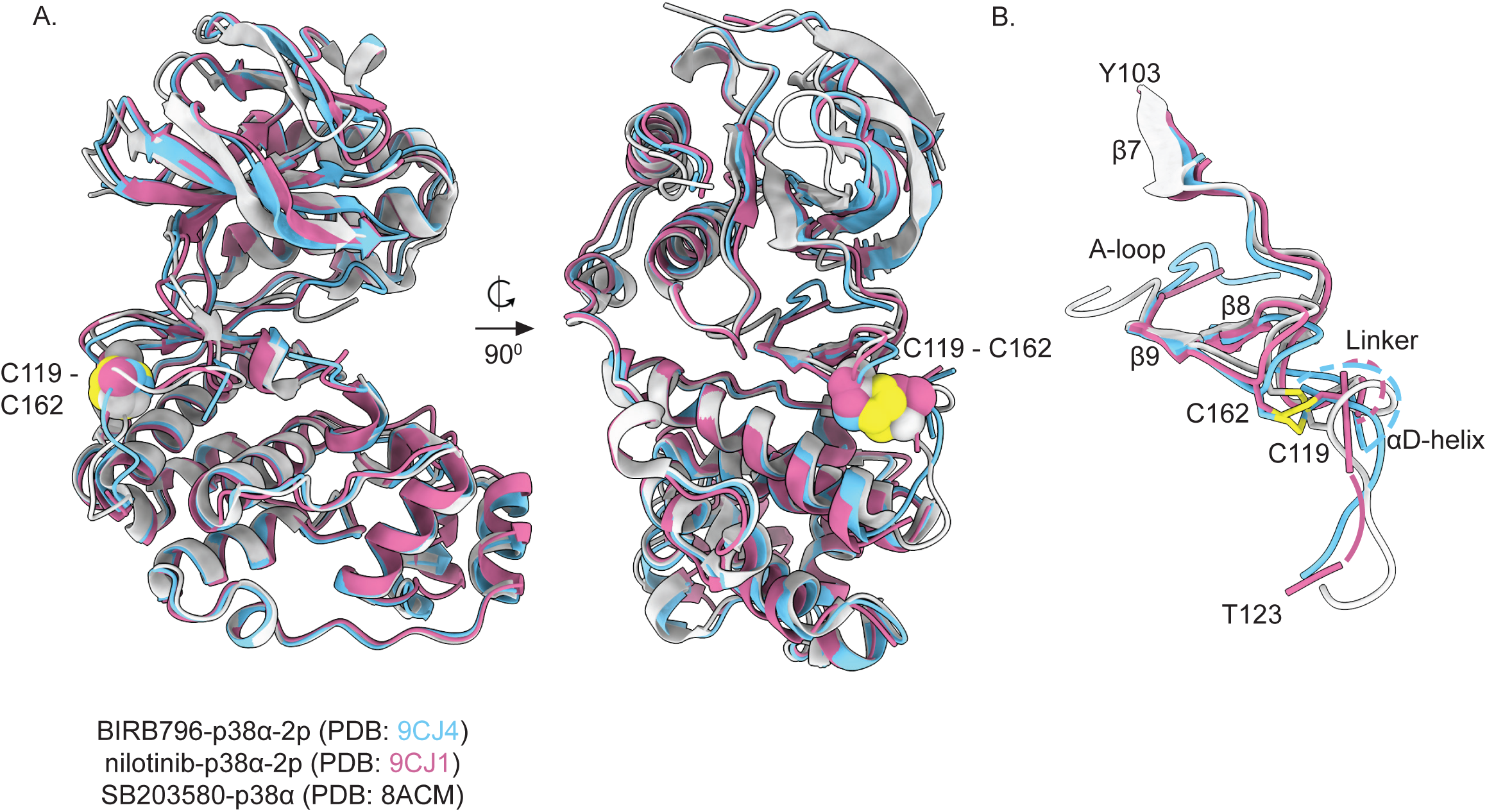
BIRB796 and nilotinib bound p38α-2p structures form a disulfide bond previously described. (A) Overlay of p38α-2p bound to BIRB796 (light blue; PDB: 9CJ4), bound to nilotinib (pink; PDB: 9CJ1), and bound to SB203580 (white; PDB: 8ACM). Disulfide bond between C119 and C162 is shown in yellow as spheres. (B) Isolated overlay of the activation loop and linker region containing the disulfide bond (yellow). In all structures, oxygen, nitrogen, phosphorous and fluorine atoms are colored red, blue, orange and light green, respectively. Crystal contacts are illustrated in Fig. S13.

**Figure S8:**
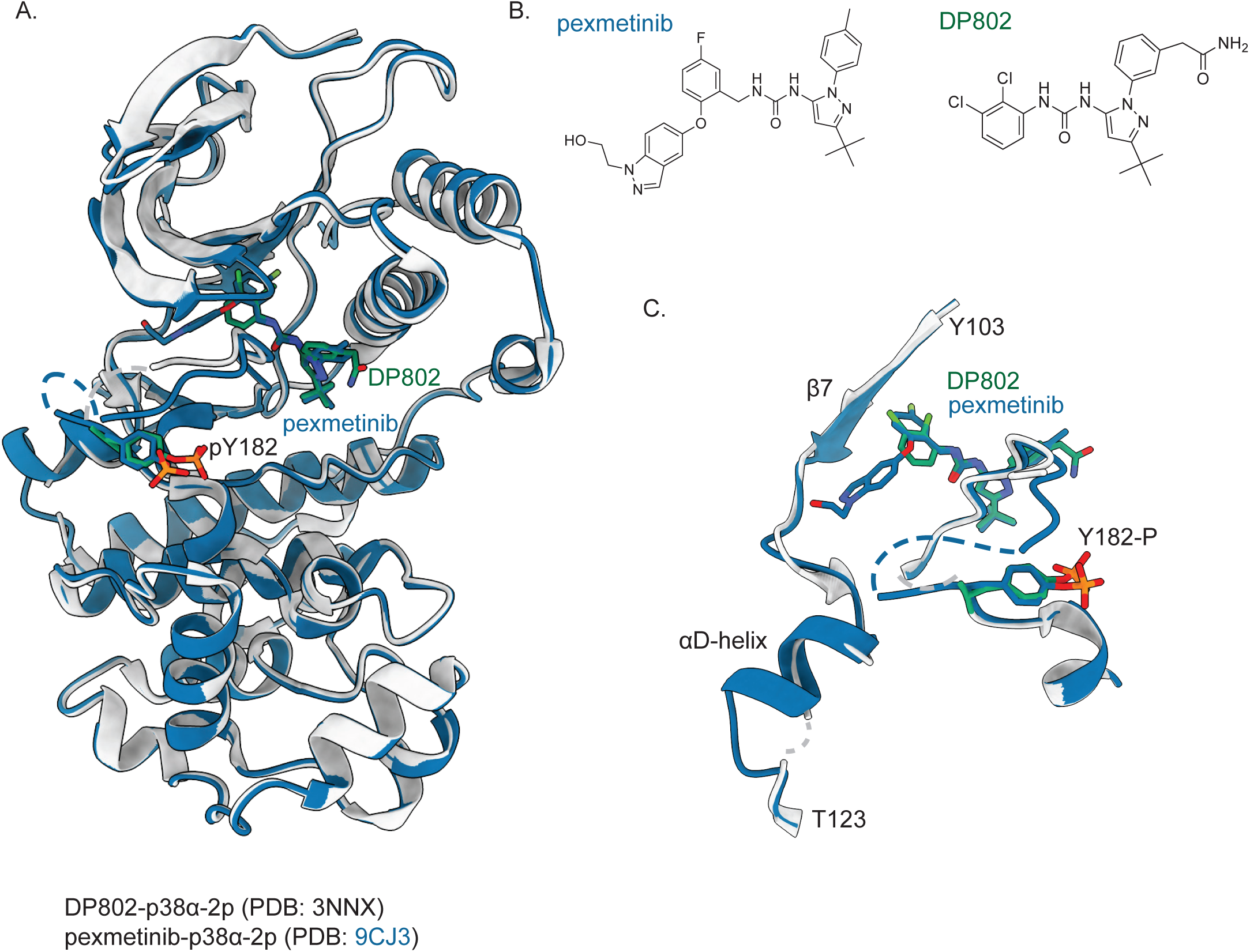
p38α activation loop flip is observed in existing phosphorylated p38α structure. (A) Overlay of X-ray crystal structures of p38α-2p bound pexmetinib (dark blue; PDB: 9CJ3) with p38α-2p bound to DP802 (white and green; PDB: 3NNX). pY182 residue in both structures is shown as sticks. (B) Chemical structures of pexmetinib and DP802. (C) Zoom in of overlayed linker and αD-helix region showing greater flexibility in the DP802-p38α-2p structure (white and green) indicated by a lack of order compared to the pexmetinib-p38α-2P (white and dark blue). In all structures, oxygen, nitrogen, phosphorous and fluorine atoms are colored red, blue, orange and light green, respectively. Crystal contacts are illustrated in Fig. S13.

**Figure S9:**
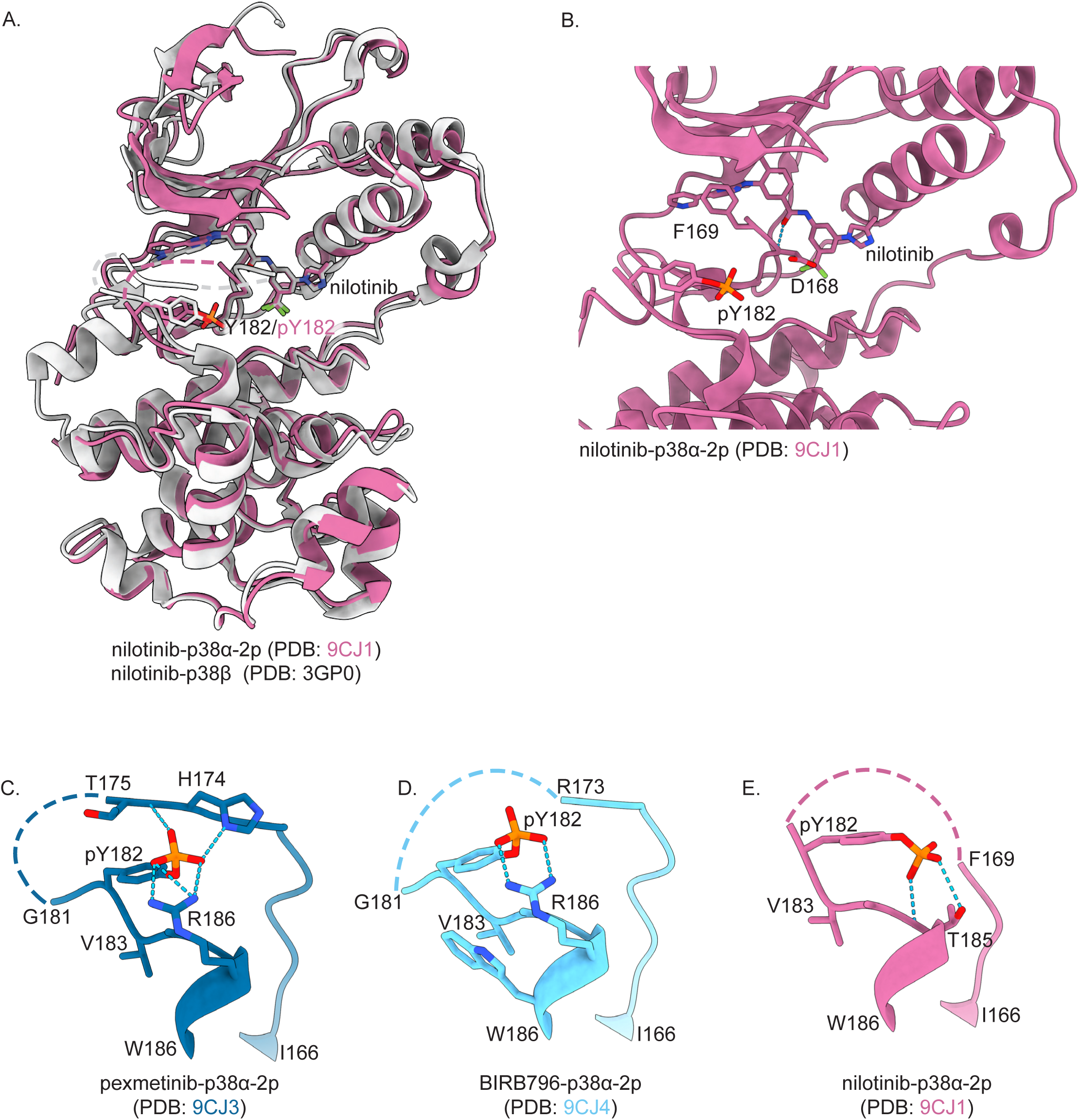
p38α activation loop flip is observed in an unphosphorylated p38β structure. (A) Overlay of X-ray crystal structures of p38α-2p bound nilotinib (pink; PDB: 9CJ1) with p38β bound nilotinib (white; PDB: 3GP0). Y182 (white) and pY182 (pink) are shown as sticks. (B) Zoomed in structure of nilotinib-p38α-2p structure (pink; PDB: 9CJ1) showing coordination of nilotinib through hydrogen bonding with D168 (blue dashed line). (C) Interaction of activation loop in p38α-2p bound to (C) pexmetinib (PDB: 9CJ3), (D) BIRB796 (PDB: 9CJ4), and (E) Nilotinib (PDB: 9CJ1). In all panels, hydrogen bonds are shown as light blue dashes. In all structures, oxygen, nitrogen, phosphorous and fluorine atoms are colored red, blue, orange and light green, respectively. Crystal contacts are illustrated in Fig. S13.

**Figure S10:**
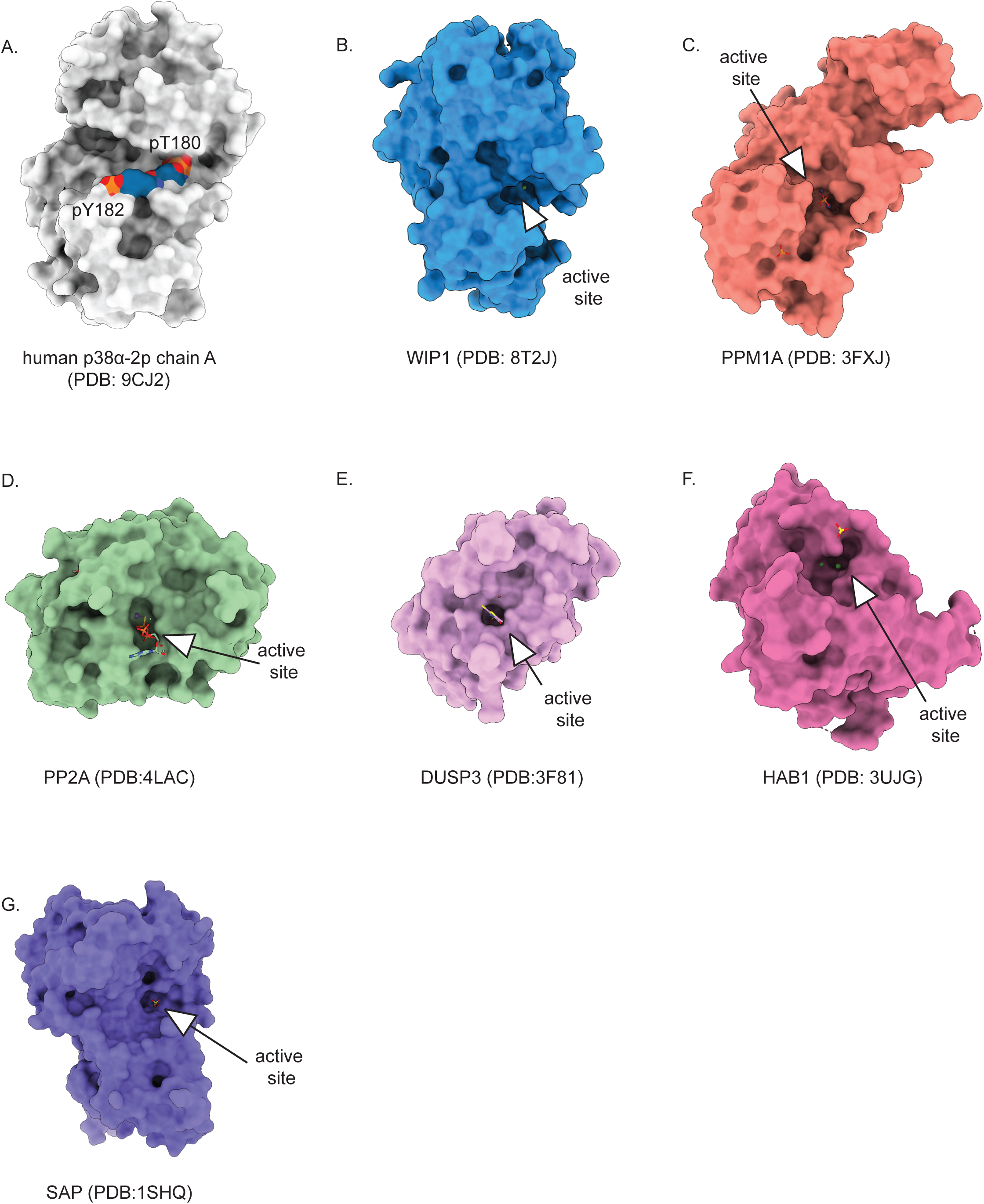
p38α-2p phosphorylation sites are inaccessible to phosphatases. (A) Surface representation of human p38α-2p chain A (PDB: 9CJ2) showing the phosphorylation sites (pT180 and pY182) are inaccessible to phosphatases. (B-G) Surface representation of various families of phosphatases that target kinases. White arrow indicating their active sites that are all inside of pockets. The active sites are occupied by the following: (B) two magnesium ions (C) phosphate ions, (D) ATPγS, (E) small molecule inhibitor SA3, (F) magnesium (green sphere) and sulfate ion (yellow sphere), and (G) a sulfate ion. In all structures, oxygen, nitrogen, phosphorous and fluorine atoms are colored red, blue, orange and light green, respectively. Crystal contacts are illustrated in Fig. S13.

**Figure S11:**
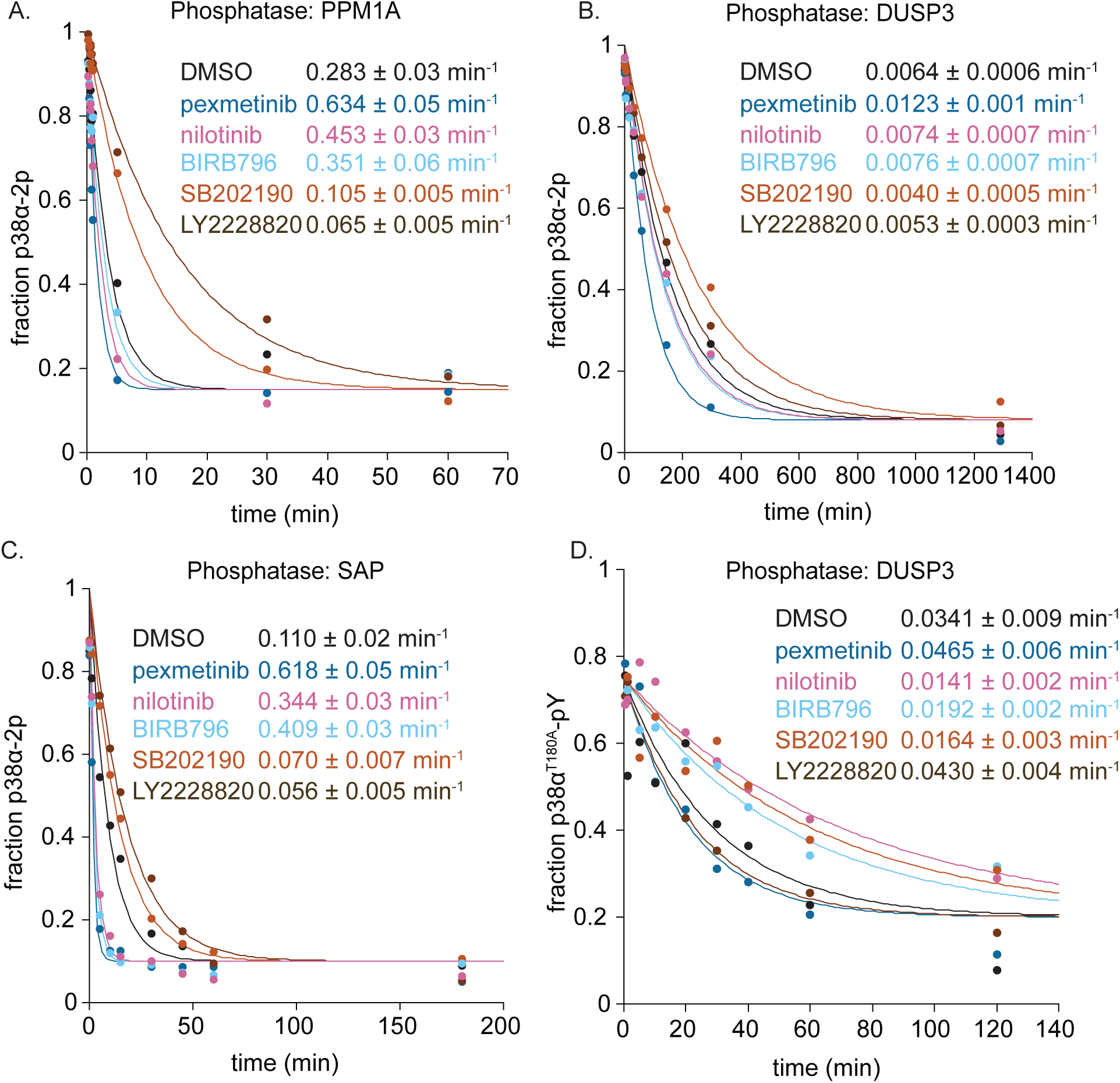
p38α dephosphorylation rates differ between phosphatases. Experimental data from which the rates were derived for the bar graphs in Fig. 5A-B. (A-C) Single turnover kinetics of p38α-2p (0.25 µM) dephosphorylation by (A) PPM1A (0.5 µ M), (B) DUSP3 (50 µM), and (C) SAP (1 µM), in the absence or presence of excess inhibitor (1.25 µM). (A) PPM1A was measured at room temperature while (B) DUSP3 and (C) SAP were measured at 37 °C to increase rate of dephosphorylation. All data is fit to an exponential decay (see methods section, n=2 ± error of the fit). (D) Single turnover kinetics of p38α^T180A^-pY (0.25 µM) dephosphorylation by DUSP3 (15 µM) in the absence or presence of excess inhibitor (1.25 µM) conducted at room temperature. All data is fit to an exponential decay (see methods section, n=2 ± error of the fit).

**Figure S12:**
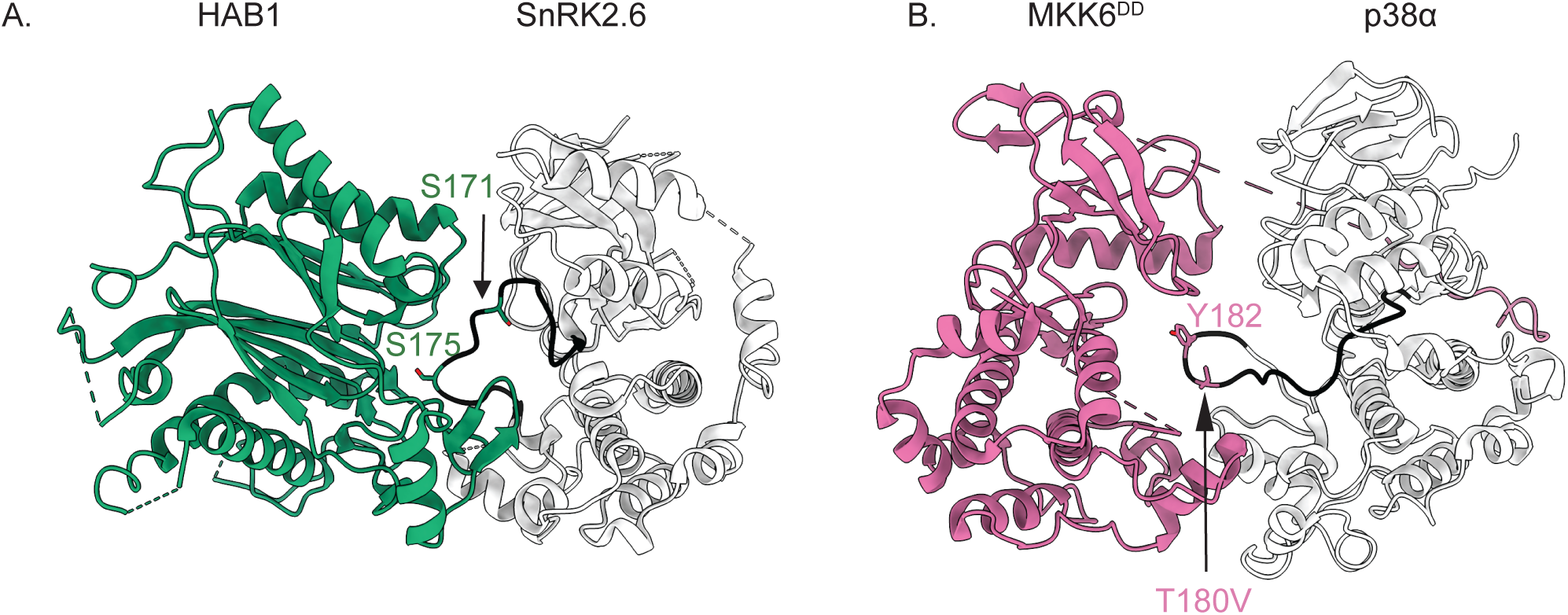
Activation loop shift is required for phosphorylation and dephosphorylation of the activation loop phospho-sites in kinases. (A) PPM plant phosphatase HAB1 (green) bound to unphosphorylated SnRK2.6 kinase (white; A-loop in black) (PDB: 3UJG). S171 and S175 of SnRK2.6 are shown as spheres and sticks in green. (B) Ser/Thr kinase MKK6^DD^ (pink) bound to unphosphorylated p38α^T180V^ (white; A-loop in black) (PDB: 8A8M). Y182 and T180V of p38α are shown as sticks in pink. In all structures, oxygen, nitrogen, phosphorous and fluorine atoms are colored red, blue, orange and light green, respectively.

**Figure S13:**
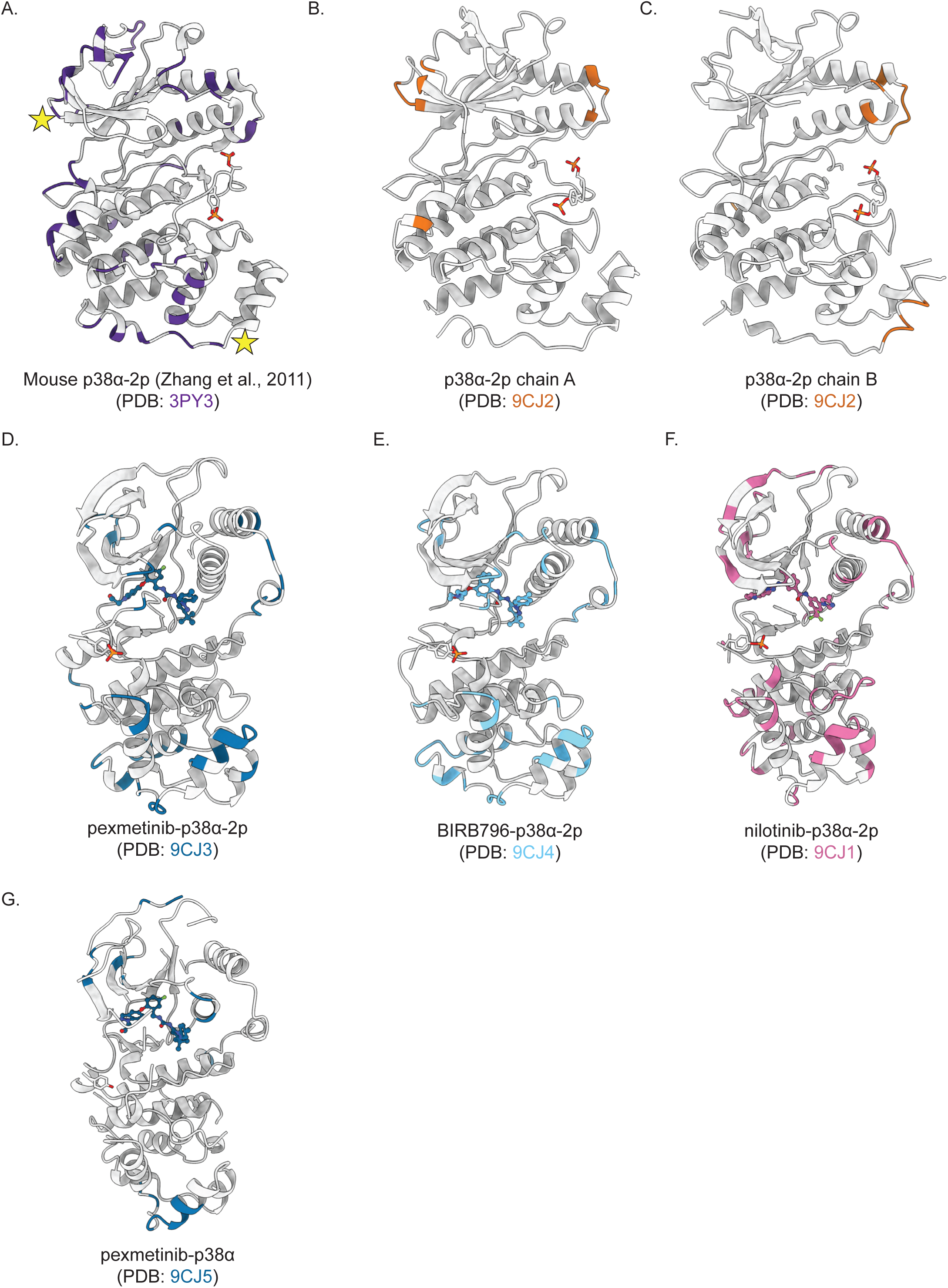
All crystal structures have distinct crystal contacts. Crystal contacts of each chain are highlighted in an associated color. Phosphorylation sites (pT180 and pY182) are shown as sticks. (A) Mouse p38α-2p (purple; PDB: 3PY3) Yellow stars indicate the two residues unique to mouse p38α compared to human p38α. H28 (top) is a crystal contact and A263 (bottom) is adjacent to a crystal contact. (B) Human p38α-2p chain A (orange; PDB: 9CJ2). (C) Human p38α-2p chain B (orange; PDB: 9CJ2). (D) Human p38α-2p bound pexmetinib (dark blue; PDB: 9CJ3). (E) Human p38α-2p bound BIRB796 (light blue; PDB: 9CJ4). (F) Human p38α-2p bound nilotinib (pink; PDB: 9CJ1). (G) Human p38α bound pexmetinib (dark blue; PDB: 9CJ5). In all structures, oxygen, nitrogen, phosphorous and fluorine atoms are colored red, blue, orange and light green, respectively.

**Table S1:**
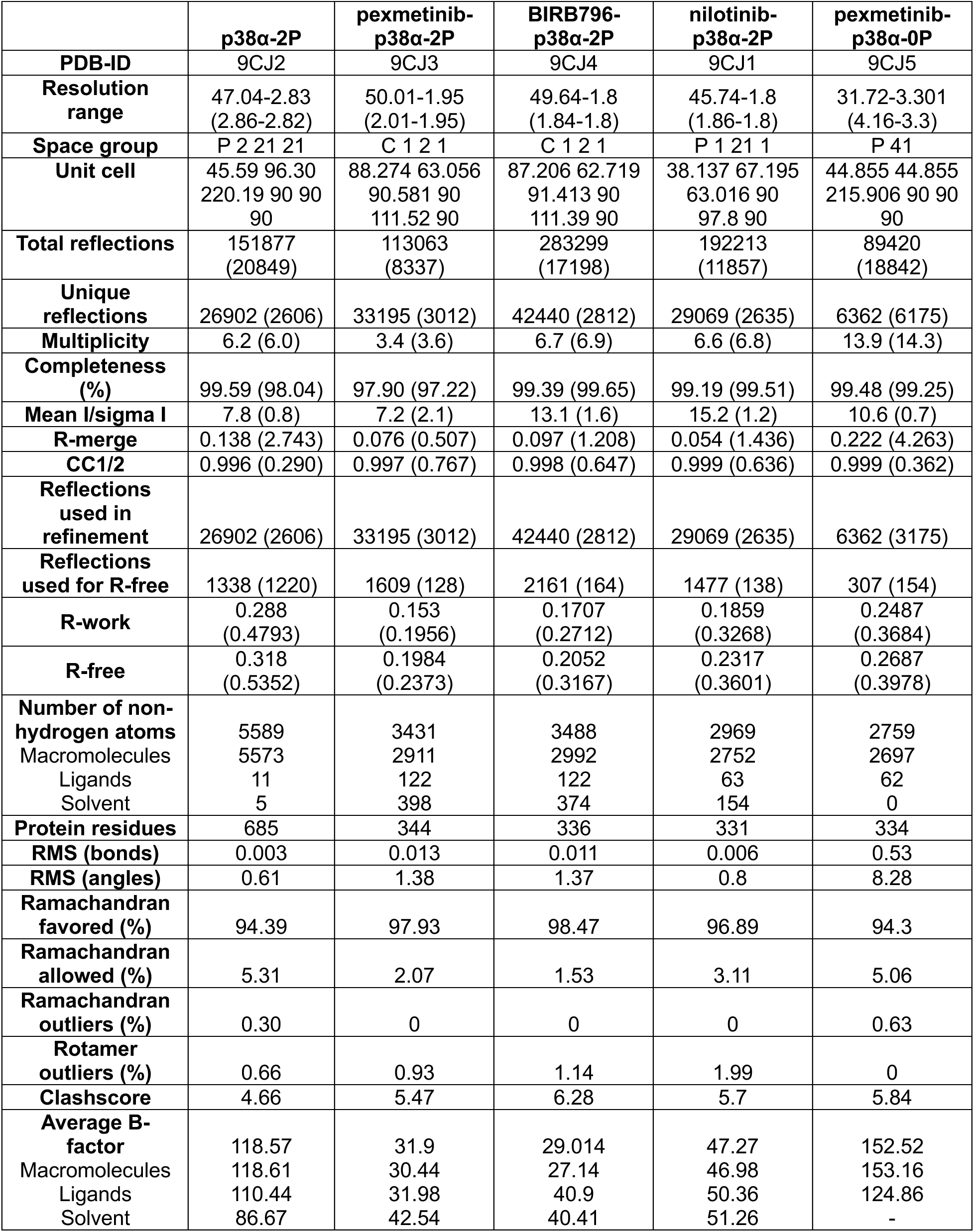
X-ray structures data collection and refinement statistics. Statistics for the highest-resolution shell are shown in parentheses.

